# A systematic review on leptospirosis in cattle: a European perspective

**DOI:** 10.1101/2023.03.07.531463

**Authors:** Cynthia Sohm, Janina Steiner, Julia Jöbstl, Thomas Wittek, Clair Firth, Romana Steinparzer, Amélie Desvars-Larrive

## Abstract

**Background:** Leptospirosis is a zoonotic disease which is globally distributed. Bovine leptospirosis often results in economic losses through its severe impact on reproduction performance. However, a clear overview of the disease characteristics in European cattle is lacking. The objective of this review was to summarise the current knowledge and state of the research on the epidemiology of bovine leptospirosis in Europe.

**Methodology:** We conducted a systematic literature review following the PRISMA guidelines. We screened four electronic databases (Pubmed, Web of Science, Scopus, CABI) and included studies published between 2001 and 2021, in English, German, and French. Identified papers were filtered according to predefined inclusion and exclusion criteria.

**Results:** Sixty-two studies were included. Reported seroprevalences were remarkably variable among studies, probably reflecting local variations but also heterogeneity in the study designs, laboratory methods, and sample sizes. The five most reported circulating serogroups in European cattle were Sejroe, Australis, Grippotyphosa, Icterohaemorrhagiae, and Pomona. Abortion and fertility disorders were the most frequently reported signs of leptospirosis in European cattle and were generally associated with chronic infections. The acute form primarily affected juveniles and foetuses. Risk factors positively associated with leptospirosis in cattle were diverse, related to environmental (e.g. geographic location), climatic (e.g. flooding), and medical (e.g. presence of other diseases) parameters, as well as farming practices (e.g. purchase policy, herd size) and individual factors (e.g. animal age and breed).

**Conclusions:** Clinical features of bovine leptospirosis in Europe cover a large range of signs and confirmation of infection requires laboratory tests. The epidemiology of the disease is very local, most probably influenced by context-specific factors. This work highlights several research gaps, including a lack of research data from several countries, a lack of methodological harmonisation, a lack of large-scale studies, an underrepresentation of beef herds in the studies, and a lack of molecular investigations.

**Author Summary:** Leptospirosis is a zoonotic disease that has been reported in cattle worldwide. This review systematically evaluated the available literature (2001-2021) on (i) the methods of diagnostics, (ii) (sero)prevalence, (iii) circulating serogroups/serovars, (iv) clinical signs, (v) and risk factors associated with leptospirosis in European cattle. We found that the prevalence of the disease was very variable among studies. Similarly, a wide range of clinical signs were described, but the most frequently reported ones were abortion and fertility disorders. Risk factors of infection in cattle included herd size, purchase of animals, access to pasture and natural water sources, contact with other animal species, presence of other diseases on farm, animal age, and occurrence of extreme weather events. This review highlights that leptospirosis should be considered as a differential diagnosis in case of abortion or reproductive failure in European cattle and emphasises the need for integrated disease prevention and control measures at farm or regional level. We identified several research gaps, particularly a lack of research data from several countries, a lack of large-scale studies, an underrepresentation of beef herds, and a limited use of molecular tools in the studies.

## Introduction

Leptospirosis is a zoonotic disease (i.e. it can affect both, humans and animals), which is distributed worldwide. It is categorised by the World Health Organization (WHO) as a neglected zoonotic disease, i.e. a disease, which burden on society is largely underestimated, such that dedicated resources to prevent or control it are very limited [1]. Costa et al. estimated that, globally, the annual number of human leptospirosis cases is as high as 1.03 million while the disease causes 58,900 deaths annually [2]. Most outbreaks occur in tropical regions, but cases are also reported from temperate areas [3] where they are often related to recreational or professional exposure to contaminated water or animals [4,5].

The causative agent, Leptospira spp., belongs to the order Spirochaetales. The bacterium is spiral-shaped, hooked on one or both ends and shows a corkscrew motility [6]. Leptospira spp. has been reported in a wide range of mammals worldwide [7]. Transmission occurs primarily through direct or indirect (i.e. via contaminated water or soil) contact with the urine of infected animals [3,8,9]. The bacteria can enter the body through the mucous membranes or damaged skin [8,9]. Following a leptospiremic phase, the bacteria colonise various organs, especially the kidneys from where they are then shed intermittently in the urine [6,9–11]. In cattle and sheep, the male and female genital tracts have been described as a major extra-renal colonisation sites of the bacteria [12–16]. Transplacental transmission may also be observed when infection occurs during pregnancy, generally resulting in abortion or stillbirth [9,17].

The phylogenetic classification organises Leptospira species based on DNA relatedness. In 2019, using large-scale whole-genome sequencing, Vincent et al. revised the classification so that the genus Leptospira spp. now contains 68 named species [18,19]. This genetic classification coexists with the historical, serological classification, which recognises more than 300 serovars of Leptospira grouped into serogroups [20,21] based on the expression of the surface-exposed lipopolysaccharide (LPS).

The gold standard for the detection of leptospiral antibodies is the microscopic agglutination test (MAT), which uses live cultures of Leptospira strains that are tested at different dilutions against patient (animal or human) serum and generally requires paired, acute and convalescent, serum samples taken at appropriate time intervals for confirmation [6]. The enzyme-linked immunosorbent assay (ELISA) is also widely used for the serological diagnostic of leptospirosis [9]. In contrast to the ELISA assays used for diagnosis of human infection, which are generally broadly reactive and can evidence a broad range of serogroups [22], few serovar-specific assays have been developed in veterinary medicine that can detect serovar-specific antibodies. For example, there is an ELISA assay to detect infection by serovar Hardjo from blood or milk sample in cattle [23]. Isolation of the bacteria via culture is also possible but is not suitable for the diagnosis of acute leptospirosis due to the fastidious growth of the bacteria, which often requires specific culture media and might take several weeks to provide a positive result [5,20]. The seroptyping of circulating Leptospira by MAT is especially important for clinical and epidemiological investigations because it may indicate which animal reservoir is involved in an infection [24]. However, the method is time-consuming and usually restricted to reference laboratories [5,25]. PCR-based strategies give faster results when compared to culture and have proven to be useful to detect Leptospira from urine, cerebrospinal fluid, or blood samples during the early stages of the disease [25] as well as in the urine, kidney, or genital tract of chronic animal carriers [12,26,27].

The environmental persistence of Leptospira and its epidemiology rely on the chronic colonisation of the kidneys of maintenance animals. Several serovars exhibit preferential association with certain hosts [28]. An animal reservoir may remain symptom-free while excreting the bacteria in its urine, either transiently or for its entire life [6,26,29]. Rodents, especially rats, are considered the main source of infection in humans and animals [30]. Although cattle are recognised as the maintenance host of serovar Hardjo (serogroup Sejroe) [28], infection of cows with this serovar is often associated with chronic infection and generally leads to abortion, fertility disorders, and decrease in milk yield [6,9,11]. In cattle, leptospirosis not only represents an occupational zoonotic disease to e.g., farm workers and veterinarians [3], but it also has an important economic impact due to the associated reproductive and non-reproductive losses to production [31,32].

Leptospirosis in cattle has been detected worldwide. Previous reviews referring to bovine leptospirosis, were conducted for Africa [33] and Latin America [34], however, to date, no work has summarised the knowledge on cattle leptospirosis from a European perspective. To characterise the epidemiology of bovine leptospirosis in Europe, we conducted a systematic literature review and examined the extent and nature of research activities pertaining to this topic over a 20-year period, 2001-2021. We addressed the following questions:

- Which laboratory methods are usually performed and on which types of samples?
- Where was bovine leptospirosis studied and what is the (sero)prevalence in the different study sites?
- Which serovars/serogroups are circulating in cattle in Europe?
- Which clinical signs are associated with leptospirosis in European cattle and what are the associated histo-pathological findings?
- What are the risk factors for Leptospira infection in European cattle?

## Methods

### Literature search strategy

From 7 June to 26 August 2021, we performed a literature search following the PRISMA (Preferred Reporting Items for Systematic Reviews and Meta-Analyses) guidelines for systematic reviews [35] (S1 PRISMA Checklist). We used the following electronic databases: Pubmed, Web of Science, Scopus, and CABI. The search terms included the following keywords “lepto*”, “cattle” “cows,” and “cow”. The keywords pertaining to the targeted species were combined using the Boolean operator “OR” and combined with the targeted disease/bacteria using “AND” (S1 Table). Additional papers were identified through internet-based search engines such as Google and Google Scholar and by hand-searches of the references cited in the reviewed studies. The period for articles to be included was set from 1 January 2001 to the date of search (26 August 2021), thereby covering over 20 years.

### Paper selection and screening

First, citation data, title and abstracts were compiled and de-duplicated in Mendeley Reference Manager and exported into Microsoft Excel 2016. Two reviewers (CS and ADL) independently screened all titles and abstracts. Titles/abstracts were selected if they contained qualitative and/or quantitative data on leptospirosis in cattle and if the study was conducted in a European country, as defined by the most common geographical definition of Europe, i.e. the land bordered by the Arctic Ocean to the north, the Atlantic Ocean to the west, the Mediterranean Sea to the south, and the Ural Mountains to the east. Case reports, outbreak description, epidemiologic surveys, reviews or epidemiological reports were included. If studied species or geographic region of the research were not specified in the title or abstract, the publication proceeded to the next selection steps.

Articles deemed eligible in the first round of screening were retrieved in full text format and reviewed independently by CS and ADL. Papers were excluded if the study was not pertaining to bovine leptospirosis, was performed outside Europe and/or was written in languages other than English, German, or French. Editorials, commentaries, book chapters, and conference proceedings were excluded. We also excluded papers with a focus on immunology, vaccine strategy, vaccine efficacy, or development of diagnostic tools and methods. Additionally, papers that did not include any epidemiological (e.g. data on prevalence, risk factors, circulating strains/serovars/serogroups) or clinical data were excluded. Disagreements between reviewers were discussed and resolved by consensus.

### Data extraction and synthesis

Two reviewers (CS and ADL) developed a data extraction sheet in Microsoft Excel 2016 and each reviewer extracted data independently. The following information was extracted from the included papers: bibliographic information (including citation, type of paper, year of publication), study purpose, study design, country and region of the study, sampling unit (animal or herd), number of sampling unit(s) investigated, sample type(s), laboratory methods, positivity threshold, study period, reported prevalence and/or incidence, reported serogroup(s) and/or serovar(s), reported genomospecies, clinical presentation, histological or necropsy findings, and production type (i.e. dairy, beef, mixed). Regarding risk factors for Leptospira infection, the following data was collected: investigated risk factor, dependent variable (i.e. assessed output), whether or not a statistical analysis was performed, direction of the association between the dependent variable and the risk factor (e.g. positive, negative, not evidenced), statistical model used, statistics reported, value of the statistics, 95% confidence interval, and p-value. Furthermore, each risk factor was attributed a category allowing to group the numerous risk factors into broader themes, leading to a better overview and more comprehensive understanding of the risk associated to Leptospira infection in European cattle. For example, icteric abortion was categorised as clinical signs; animal introduction (e.g. purchase) was categorised as biosecurity. All risk factors and their assigned risk categories are shown in S1 Appendix. Data extracted by one researcher was double checked by the other. Two additional independent reviewers (JS and JJ) performed a final curation and validation of the extracted data. Summary statistics, tables and figures were computed in R v.4.0.3 [36] using RStudio [37].

## Results

### Selected studies

The PRISMA flow chart presented in Fig 1 summarises the search strategy. The database search retrieved 2,181 articles (570 from Pubmed, 926 from Scopus, 537 from Web of Science, and 148 from CABI). Three articles were manually identified through other sources and included. After exclusion of duplicates (n = 1,128), the title and abstract of 1,056 records were screened. Of these, 898 articles were excluded based on our exclusion criteria: i.e. the article was not located in Europe (n = 435), was not about Leptospira spp. or leptospirosis (n = 195), did not involve cattle (n = 67), did not contain epidemiological data (n = 67), dealt with immunology (n = 22), vaccine efficacy (n =37), or diagnostic tools or methods (n = 50). Twenty-five conference papers and posters were also excluded. The remaining articles (n = 158) were subjected to full-text screening. The same exclusion criteria were applied. Many studies had to be excluded on the basis of more than one criterion; numbers given in Fig 1 indicate the primary criterion of exclusion that was identified. A total of 62 articles were ultimately included in the systematic review. The details of the included studies are shown in S1 Appendix.

**Fig 1.**
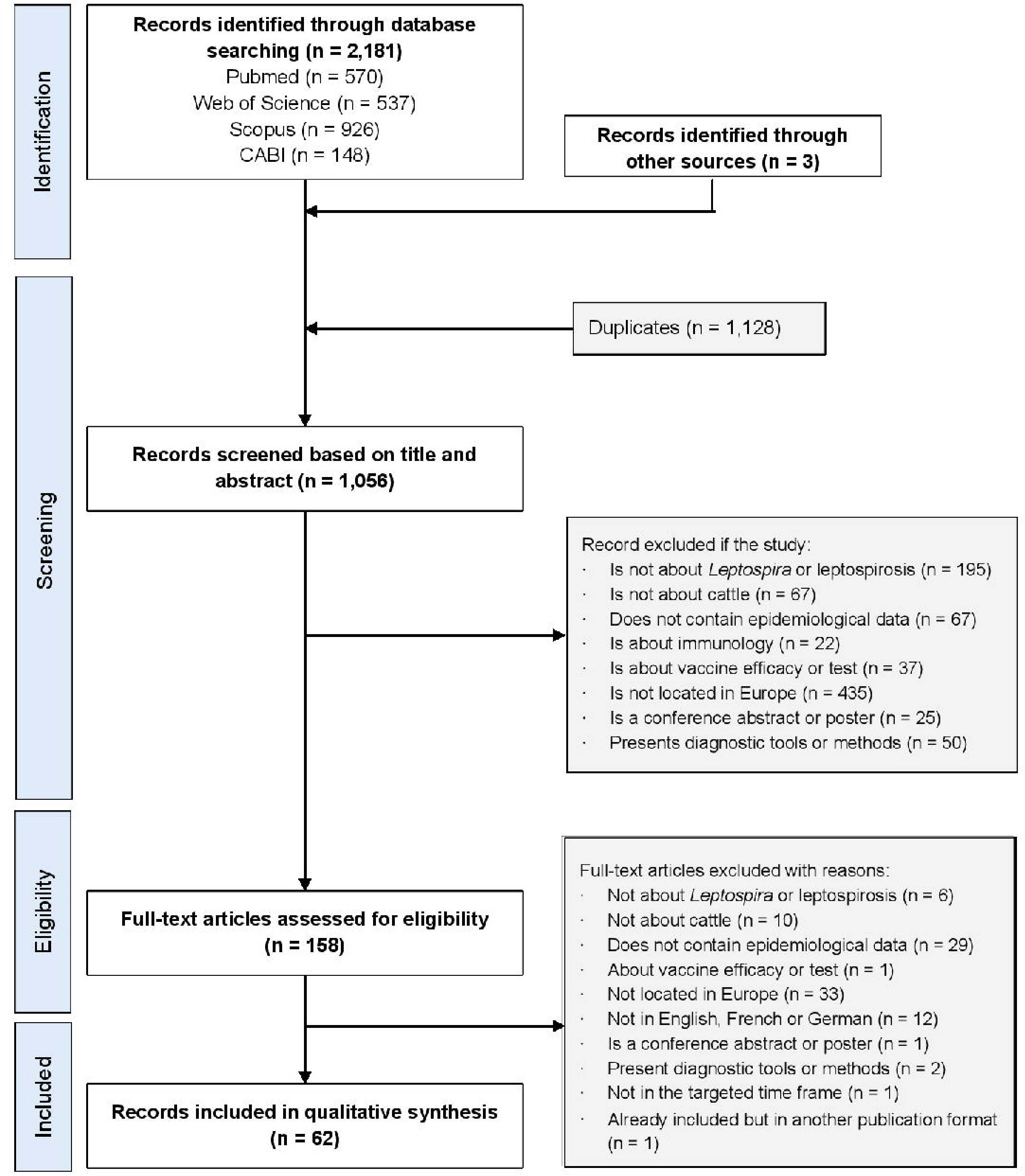
Identification and selection of the included papers according to the PRISMA guidelines.

Included articles, published between 2001 and 2021 (Fig 2), covered studies conducted in 18 different European countries (Fig 3). The geographic distribution of the included studies showed spatial heterogeneity, with the highest number of studies conducted in the United Kingdom (n = 13, 21%) [38–50], the Republic of Ireland (n = 8, 12.9%) [51–58], and France (n = 8, 12.9%) [59–66]. Fifty articles were classified as research papers (80.6%), nine were regional or national disease surveillance or laboratory reports (14.5%), and three were articles published in professional veterinary journals (4.8%). Regarding study design, four were case-control studies (6.5%), 16 were clinical case investigations (25.8%), two were cohort study (3.2%), 36 were cross-sectional studies (58.1%), and seven were longitudinal studies (11.3%) (three studies contained more than one research design).

**Fig 2.**
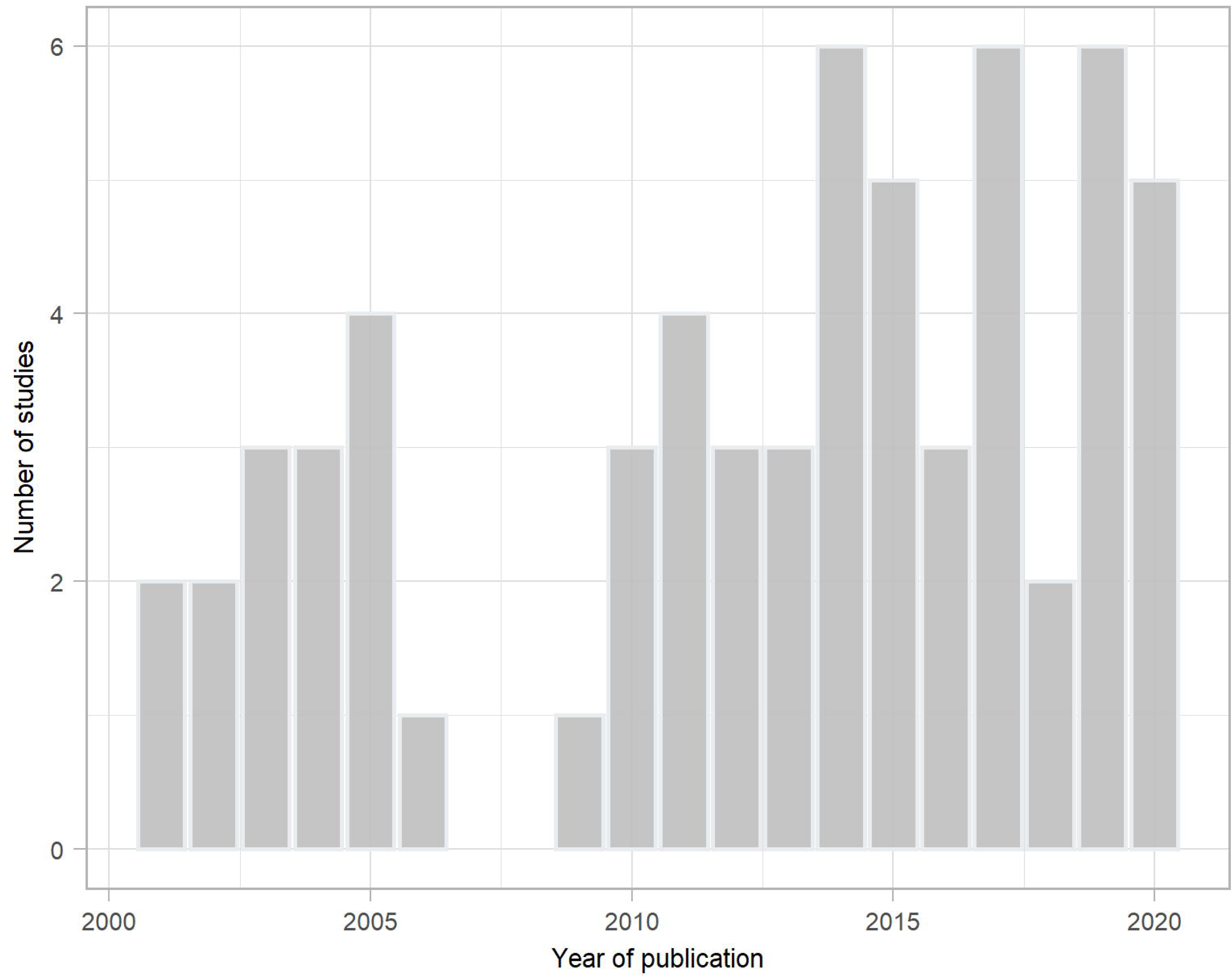
Bar chart of the distribution of the year of publication of the 62 studies included in the systematic review on cattle leptospirosis in Europe, 2001-2021.

**Fig 3.**
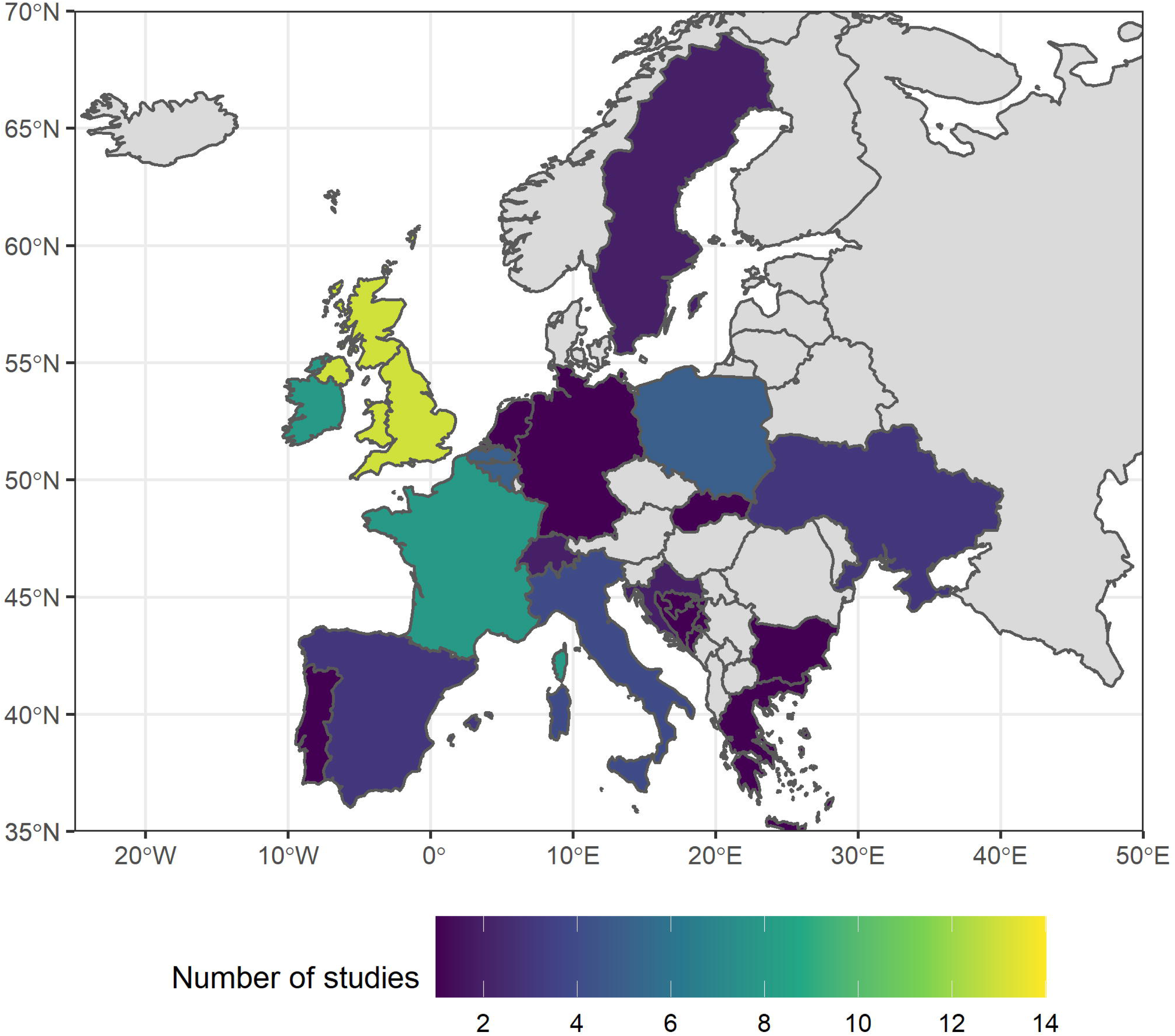
Number of studies per country pertaining to bovine leptospirosis published in Europe between 2001 and 2021 and selected for this systematic review. Grey: no studies included. The map was produced using R packages. The map is from the package "rnaturalearth" and is licensed-free from South, A. (2017). rnaturalearth: World Map Data from Natural Earth, R package version 0.1.0 (https://CRAN.R-project.org/package=rnaturalearth).

The production type was mentioned in 34 articles (54.8%). A majority of the studies examined dairy cattle (n = 23, 37.1%) [38,44,45,47–50,52–54,58,64,65,67–76] whereas beef cattle were occasionally considered (n = 7, 11.3%) [46,51,55,56,63,66,77]; three studies investigated both production types (4.8%) [78–80] and one study (1.6%) surveyed only female cattle, independent of the type of production [60].

### Laboratory methods

Fifty-six articles (90.3%) specified the laboratory methods used to investigate occurrence of Leptospira in cattle. Serological tests, detecting the presence of antibodies against Leptospira spp. were predominantly used. The MAT was the most commonly employed method (n = 36 studies, 58.1%) and predominantly performed on serum samples [57,59–68,71–79,81–95], although Grégoire et al. used pleural fluids from aborted foetuses as an alternative [96]. ELISA was also widely used (n = 19, 30.6%), on serum [46,51,55,56,80,85,97] or milk [45,47,48,50,52–54,69,70] samples or both [44,49,98]. Generally, the circulating serovars/serogroups, confirmed by antibody detection, were reported (in 34/36 and 16/19 studies, respectively), although the ELISA generally detects specific antibodies against serovar Hardjo only. The agglutinin-absorption test (AAT) was also used for antibody detection (n = 1) [68]. The cut-off value (i.e. the threshold that defines whether a test result is positive or negative) for the MAT was provided in 31/36 (86.1%) studies and was highly variable, ranging between 1:5 and 1:1,000 [57,59–63,66–68,71,73–79,82–92,94,96,98]. Similarly, the number of serovars included in the MAT panel (mentioned in 28 studies) showed large disparities (min. = 1; max. = 24 strains), with a median of 10.5 strains per panel [57,59–62,65–67,71,72,74,75,77–79,81–85,87–92,94,95].

PCR was occasionally performed (n = 11 studies, 17.7%) for the direct detection of Leptospira spp., e.g. in aborted foetuses (spleen, kidney, liver, brain, abomasal content), stillborn calves (spleen, liver, lung, small intestine kidney, adrenal gland, heart, brain, blood), adult cows (placenta, urine, blood) [63,66,72,74,75,77,79,85,88,96], as well as air samples [97]. Culture of Leptospira spp. (n = 3 studies, 4.8%) was performed from urine, placental cotyledons, and water samples [63,77,80], although all of them were unsuccessful. Immunofluorescence assays (n = 2) [43,88] and histology (n = 1) [88] were also used to confirm the presence of the bacteria.

Molecular methods to characterise the circulating genomospecies were rarely performed; they included multi-locus sequence typing (MLST, n = 2) [85,99], high-resolution melting analysis (HRMA) [96] and sequencing of the lfb1 gene (n = 1) [96].

In most papers a single diagnostic method was used, however, Delooz et al. combined four different methods (ELISA, MAT, PCR and MLST) [85].

### Occurrence of Leptospira infection

To calculate epidemiological parameters, i.e. (sero)prevalence, incidence, or risk, 44 studies considered the individual animal as the epidemiological unit [38–44,51–54,57,59,63–66,68,71–77,80–96,98,99] whereas 14 considered the herd (i.e. a herd was defined as positive for leptospirosis if at least one animal within this herd was tested positive) [45–48,51–55,58,61,62,69,70]; eight studies described epidemiological measures at both levels [49,50,56,60,67,78,79,97]. The seroprevalence of leptospirosis in European cattle varied greatly between countries and studies (Table 1). Overall, Belgium, France, Italy, Republic of Ireland, Spain, Ukraine, and the United Kingdom reported relative high seroprevalences of leptospirosis in cattle whereas Croatia, Germany, Greece, Poland, and Switzerland reported intermediate seroprevalences. Bosnia and Herzegovina, Bulgaria, the Netherlands, and Sweden reported relatively low seroprevalences. Within countries, regional or local estimations of the seroprevalence exhibited great variations, e.g. in Belgium [75,80,85,86,96], France [59,60,63–66], Spain [67,71,78], Ukraine [81,92,95] and the United Kingdom [44–50].

**Table 1.** Leptospira occurrence in cattle in Europe, 2001-2021. Seroprevalence: detection of antibodies through serological methods (ELISA, MAT, AAT), nce: detection of the presence of Leptospira by PCR, immunofluorescence assay, or culture. Seroprevalences and prevalences are given all laboratory s combined, respectively.

| Country | Epidemiological unit<sup>a</sup> | Seroprevalence range (No. of studies) | Prevalence range (No. of studies) | Type(s) of production system investigated (No. of studies) | References |
| --- | --- | --- | --- | --- | --- |
| Belgium | Animal | 0% – 95% (5) | 3.7% – 80.8% (2) | Dairy (1), dairy and beef (1), NA (3) | [75,80,85,86,96] |
| Bosnia and Herzegovina | Animal | 1.6% (1) | NA | Dairy (1) | [76] |
| Bulgaria | Animal | 0.4% (1) | NA | NA (1) | [94] |
| Croatia | Animal | 1% – 22.6% (2) | NA | NA (2) | [90,93] |
| France | Animal | 11.9% – 100% (6) | NA | Dairy (2), beef (2), dairy, beef and mixed (1), NA (1) | [59,60,63–66] |
| France | Herd | 38% – 100% (2) | NA | Dairy, beef and mixed (1), NA (1) | [60,61] |
| **Germany** | Animal | 1.8% (1) | NA | Dairy and beef (1) | [79] |
| **Germany** | Herd | 3.4% – 14.9% (1) | 0.7% (1) | Dairy and beef (1) | [79] |
| **Greece** | Animal | 12.6% (1) | NA | NA (1) | [83] |
| **Italy** | Animal | 0.5% – 38.7% (4) | 50% – 100% (1) | Beef (1), NA (3) | [77,82,84,89] |
| **Netherlands** | Herd | 1.1% (1) | NA | Dairy (1) | [69] |
| **Poland** | Animal | 0% – 26.8% (4) | 0% (2) | Dairy (1), NA (3) | [72,87,91,97] |
| **Poland** | Herd | 3.2% (1) | 0% (1) | Dairy (1), NA (1) | [70,97] |
| **Republic of Ireland** | Animal | 30.1% – 55.1% (2) | NA | Beef (1), NA (1) | [56,57] |
| **Republic of Ireland** | Herd | 75.9% – 93.3% (5) | NA | Beef (3), Dairy (2) | [51–53,55,56] |
| **Slovakia** | Animal | 52 positive cattle<br>(sample size unknown) | NA | Dairy (1) | [68] |
| **Spain** | Animal | 6.4% – 43.3% (3) | NA | Dairy (2), dairy and beef (1) | [67,71,78] |
| **Spain** | Herd | 36.2% – 100% (2) | NA | Dairy (1), Dairy and beef (1) | [67,78] |
| **Sweden** | Animal | 0% – 0.98% (2) | NA | Dairy (1), NA (1) | [73,98] |
| **Switzerland** | Animal | 21.4% (1) | 1.8% – 12.5% (2) | Dairy (1), NA (1) | [74,88] |
| **Ukraine** | Animal | 0.19% – 64.6% (3) | NA | NA (3) | [81,92,95] |
| **United Kingdom** | Animal | 13.7% – 41.7% (3) | 4.3% – 100% (6) | Dairy (4), NA (6) | [38–44,49,50] |
| **United Kingdom** | Herd | 21.9% – 80% (6) | NA | Dairy (5), beef (1) | [45–50] |

<sup>a</sup> Epidemiological unit investigated, e.g. if herd, the prevalence gives the number of herds positive divided by the total number of herds investigated (no information on the individual animals).

### Diversity and geographic distribution of Leptospira serovars, serogroups, and genomospecies in cattle in Europe

Overall, 53 studies reported 27 different circulating serovars, belonging to 18 serogroups, in cattle in Europe during the period 2001-2021, determined by MAT and ELISA (three studies did not specify the method) (Table 2). The five most reported Leptospira serogroups in European cattle identified by MAT were Sejroe (n = 31 studies), Australis (n =22), Grippotyphosa (n =22), Icterohaemorrhagiae (n =20), and Pomona (n =14). The most frequently reported serovars circulating in European cattle were: Hardjo (n = 21), Grippotyphosa (n = 13), Bratislava (n = 12), Pomona (n = 12), Canicola (n = 10), Icterohaemorrhagiae (n = 10), Australis (n = 7), Tarassovi (n = 7), Copenhageni (n = 6), Ballum (n = 3), Hebdomadis (n = 3), and Sejroe (n = 3). The highest serogroup diversity was found in Western Europe (15 different serogroups in 15 studies). Detailed data on serogroup/serovar information from each publication can be found in S1 Appendix.

**Table 2.**
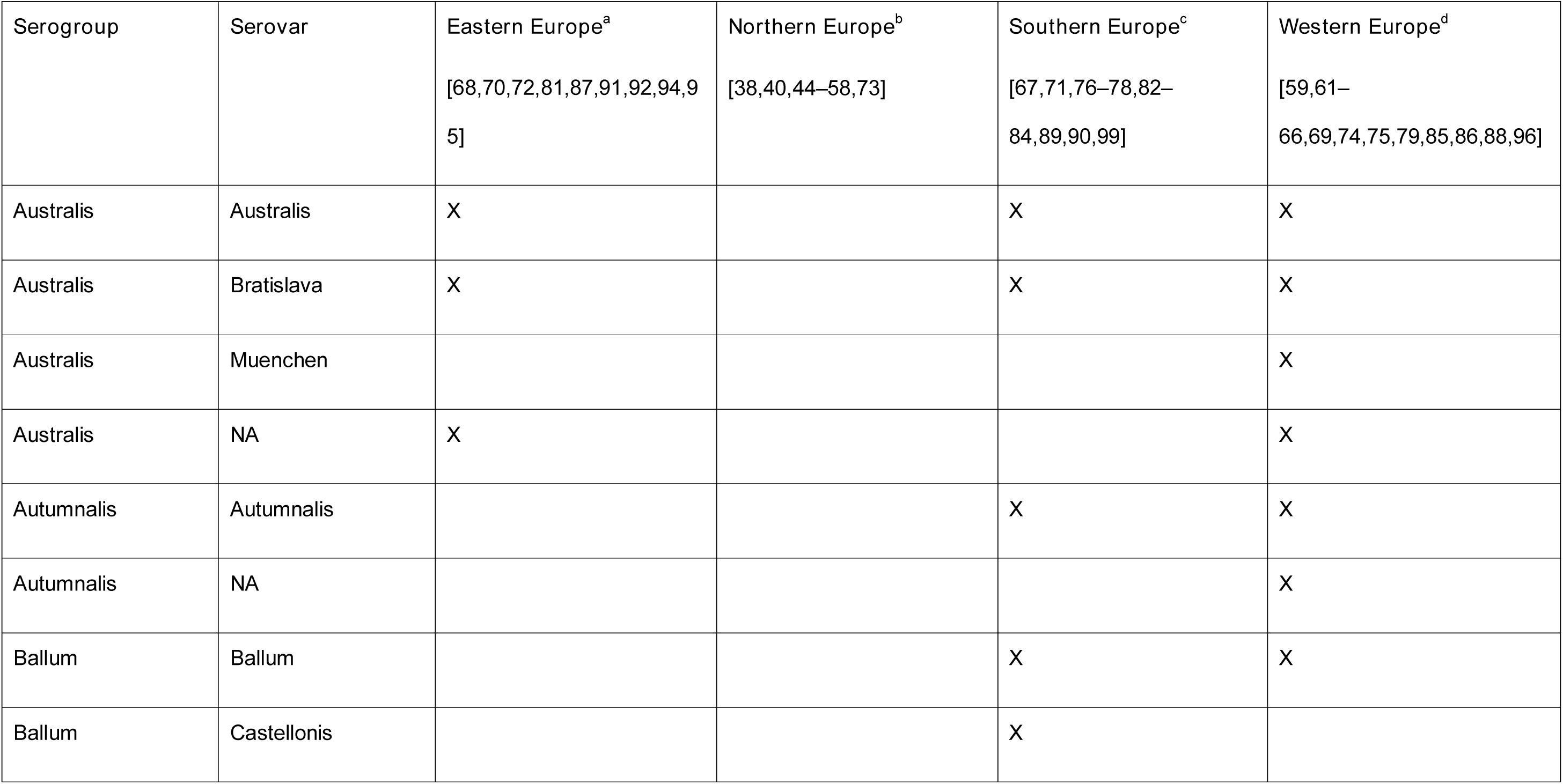
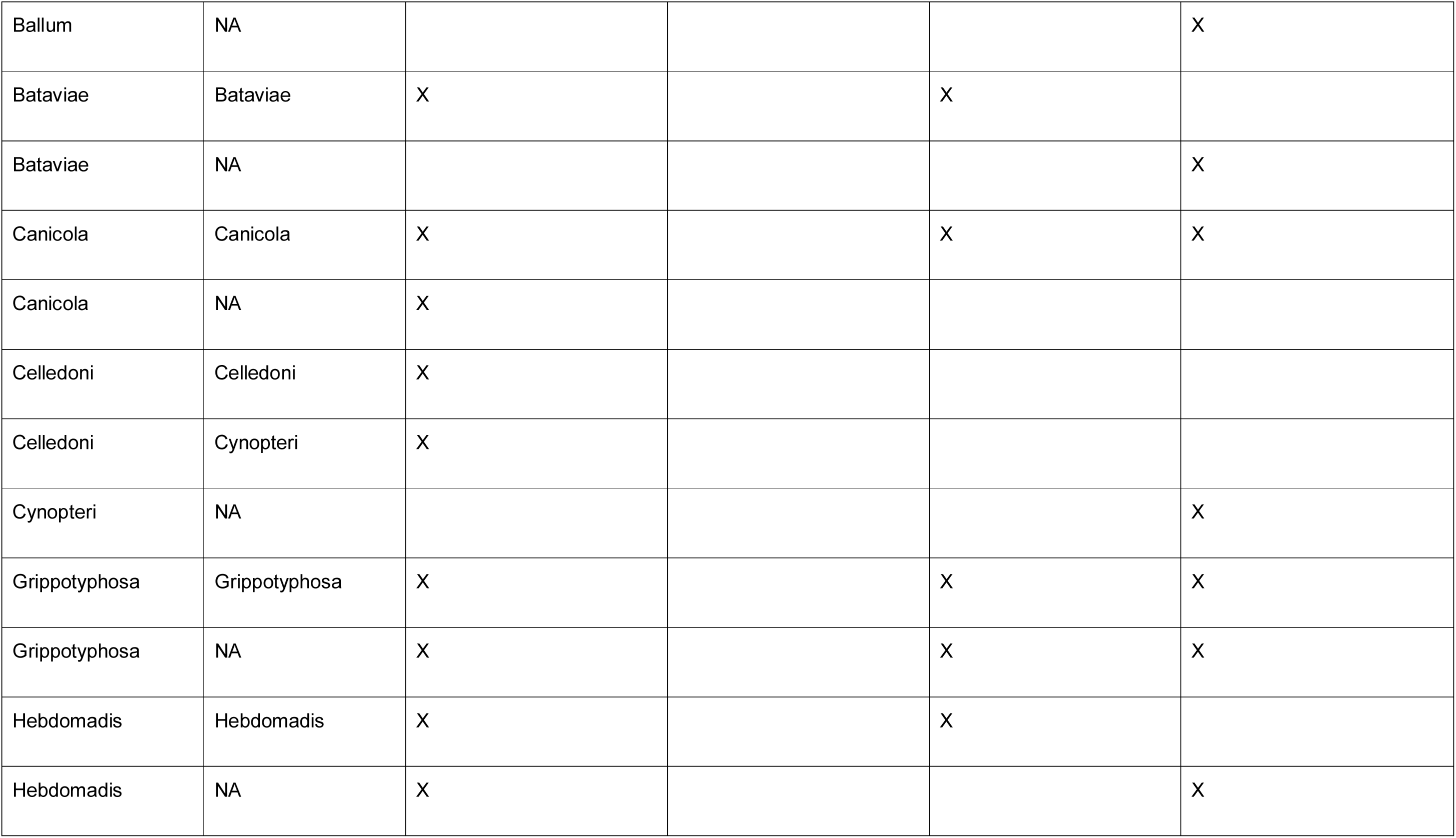
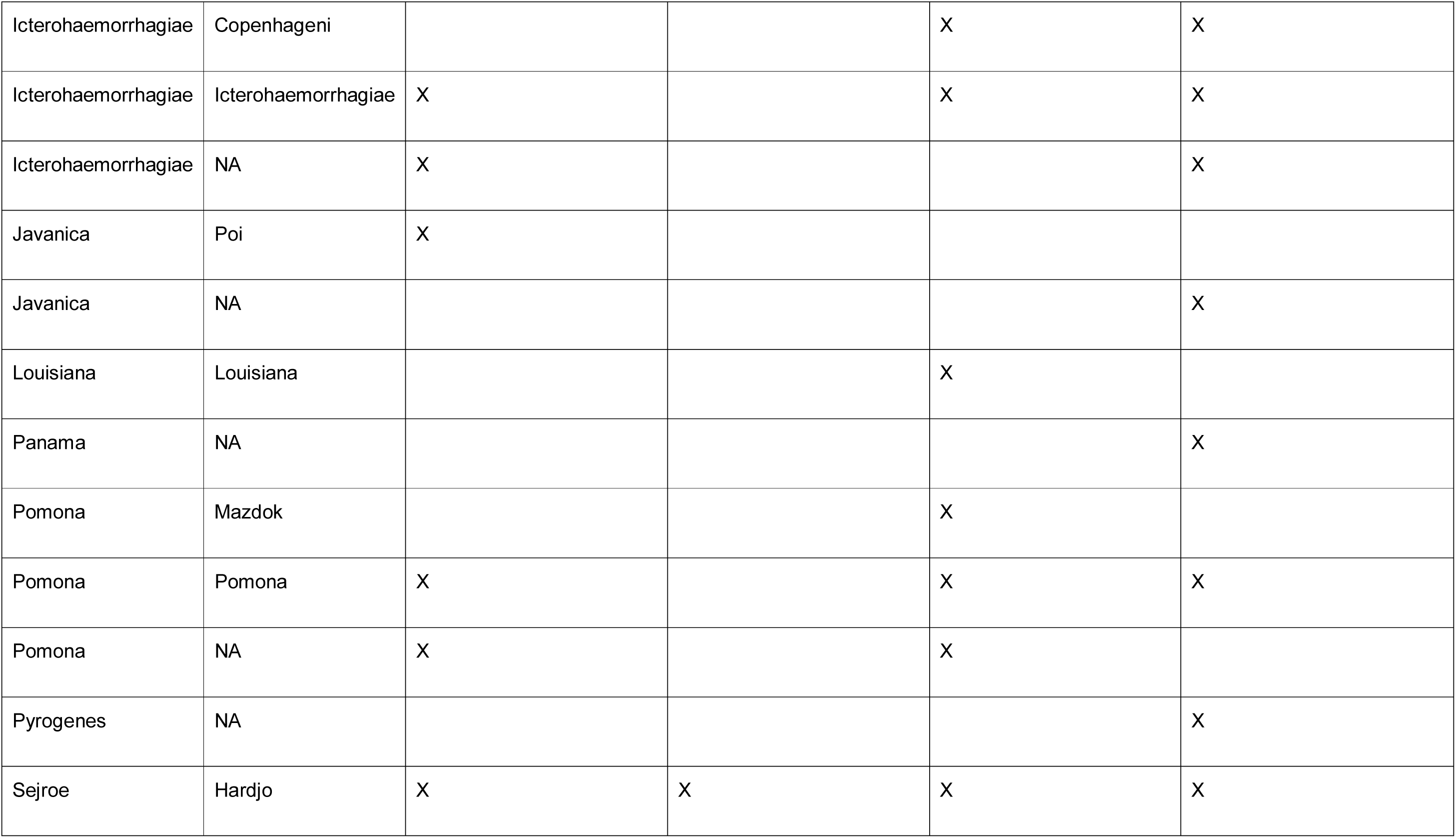
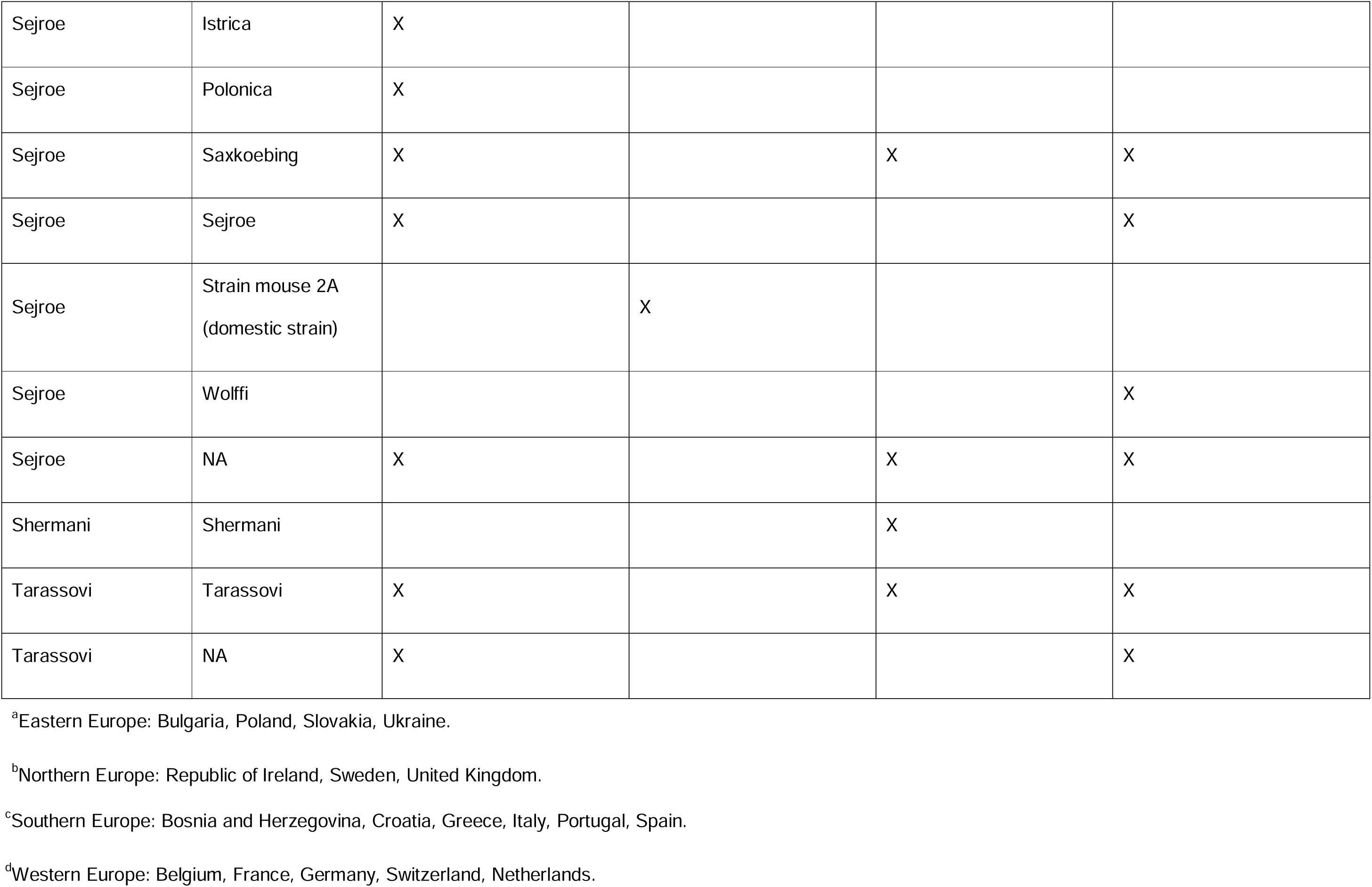
Serovars and serogroups reported in cattle in Europe, 2001-2021. We used a regionalisation of Europe as provided by UNStats [100]: Results are provided at serogroup or serovar level, depending on the information available in the study. For each serovar, when not provided, the serogroup was retrieved from the literature [101].

| Serogroup | Serovar | Eastern Europe<sup>a</sup><br>[68,70,72,81,87,91,92,94,95] | Northern Europe<sup>b</sup><br>[38,40,44–58,73] | Southern Europe<sup>c</sup><br>[67,71,76–78,82–84,89,90,99] | Western Europe<sup>d</sup><br>[59,61–66,69,74,75,79,85,86,88,96] |
| --- | --- | --- | --- | --- | --- |
| Australis | Australis | X |  | X | X |
| Australis | Bratislava | X |  | X | X |
| Australis | Muenchen |  |  |  | X |
| Australis | NA | X |  |  | X |
| Autumnalis | Autumnalis |  |  | X | X |
| Autumnalis | NA |  |  |  | X |
| Ballum | Ballum |  |  | X | X |
| Ballum | Castellonis |  |  | X |  |
| Ballum | NA |  |  |  | X |
| Bataviae | Bataviae | X |  | X |  |
| Bataviae | NA |  |  |  | X |
| Canicola | Canicola | X |  | X | X |
| Canicola | NA | X |  |  |  |
| Celledoni | Celledoni | X |  |  |  |
| Celledoni | Cynopteri | X |  |  |  |
| Cynopteri | NA |  |  |  | X |
| Grippotyphosa | Grippotyphosa | X |  | X | X |
| Grippotyphosa | NA | X |  | X | X |
| Hebdomadis | Hebdomadis | X |  | X |  |
| Hebdomadis | NA | X |  |  | X |
| Icterohaemorrhagiae | Copenhageni |  |  | X | X |
| Icterohaemorrhagiae | Icterohaemorrhagiae | X |  | X | X |
| Icterohaemorrhagiae | NA | X |  |  | X |
| Javanica | Poi | X |  |  |  |
| Javanica | NA |  |  |  | X |
| Louisiana | Louisiana |  |  | X |  |
| Panama | NA |  |  |  | X |
| Pomona | Mazdok |  |  | X |  |
| Pomona | Pomona | X |  | X | X |
| Pomona | NA | X |  | X |  |
| Pyrogenes | NA |  |  |  | X |
| Sejroe | Hardjo | X | X | X | X |
| Sejroe | Istrica | X |  |  |  |
| Sejroe | Polonica | X |  |  |  |
| Sejroe | Saxkoebing | X |  | X | X |
| Sejroe | Sejroe | X |  |  | X |
| Sejroe | Strain mouse 2A<br>(domestic strain) |  | X |  |  |
| Sejroe | Wolffi |  |  |  | X |
| Sejroe | NA | X |  | X | X |
| Shermani | Shermani |  |  | X |  |
| Tarassovi | Tarassovi | X |  | X | X |
| Tarassovi | NA | X |  |  | X |

<sup>a</sup>Eastern Europe: Bulgaria, Poland, Slovakia, Ukraine.
<sup>b</sup>Northern Europe: Republic of Ireland, Sweden, United Kingdom.
<sup>c</sup>Southern Europe: Bosnia and Herzegovina, Croatia, Greece, Italy, Portugal, Spain.
<sup>d</sup>Western Europe: Belgium, France, Germany, Switzerland, Netherlands.

The serogroup diversity is relatively stable over the 20 years (Fig 4). Three serogroups were sporadically reported, namely Cynopteri and Panama in 2018 only [85] and Pyrogenes in 2018 [85] and 2020 [96], even though these serogroups were tested in other studies [59,61,63,66,68,75,81,87,91].

**Fig 4.**
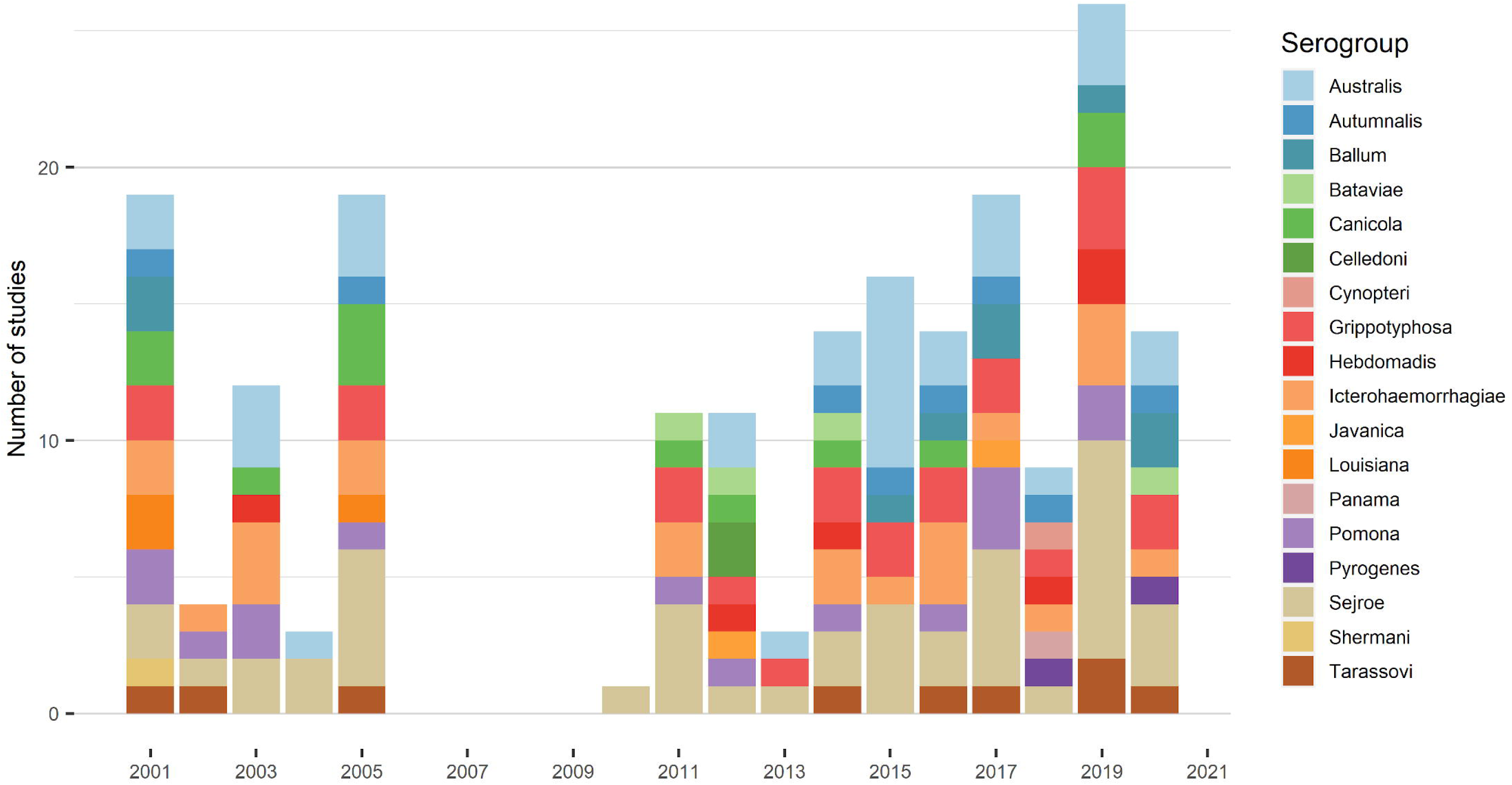
Number of studies per year reporting the different circulating serogroups in European cattle investigated through MAT, 2001-2021.

Three genomospecies were characterised in European cattle over the study period: Leptospira interrogans in Belgium, Portugal, and Republic of Ireland [58,85,96,99], Leptospira kirschneri in Belgium [85,96]. and Leptospira borgpetersenii in Portugal [99].

### Clinical signs and histo-pathological findings associated with bovine leptospirosis in Europe

Twenty-nine out of the 62 selected studies (46.8%), conducted in 13 different European countries, described clinical signs and/or histo-pathological findings associated with Leptospira infection in cattle. Clinical signs of leptospirosis differed largely with respect to the age of the animal (Table 3). Acute, serious disease rarely occurred in adult cattle and usually affected calves and foetuses [43,65,66]. Fig 5 displays the clinical signs reported in each country.

**Fig 5.**
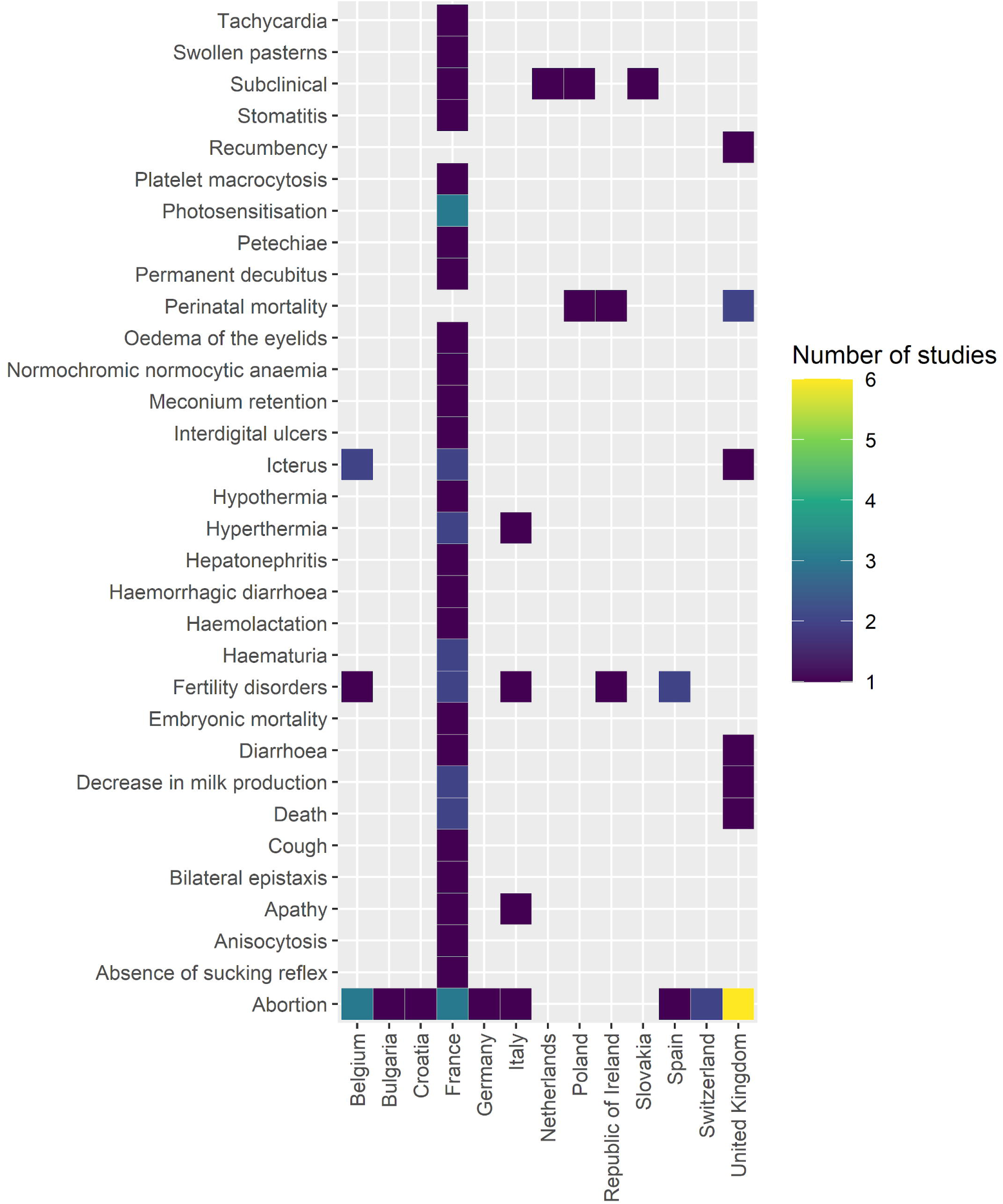
Heat map displaying the number of studies per country, reporting at least one clinical sign associated with bovine leptospirosis, Europe, 2001-2021.

**Table 3.**
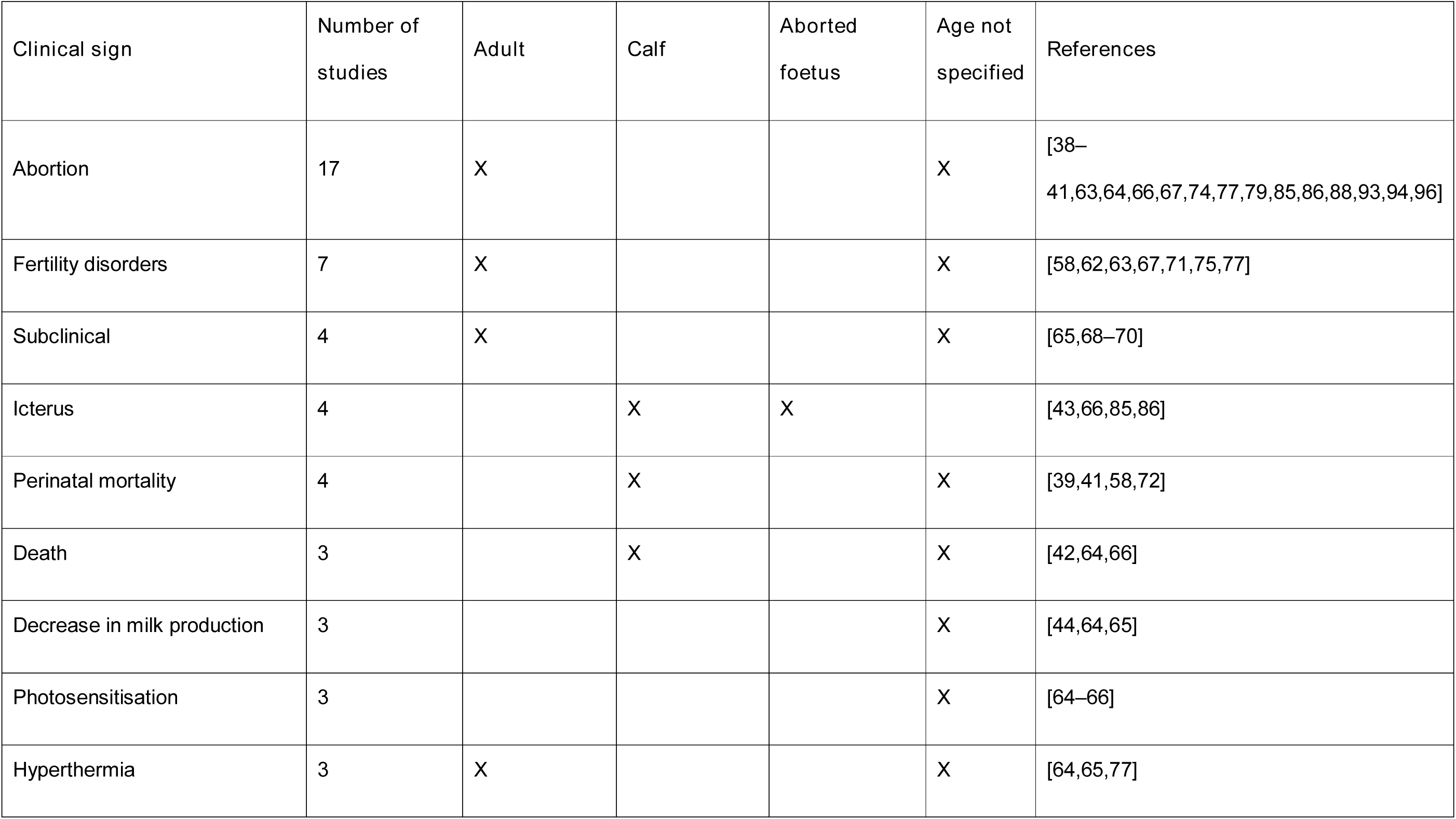
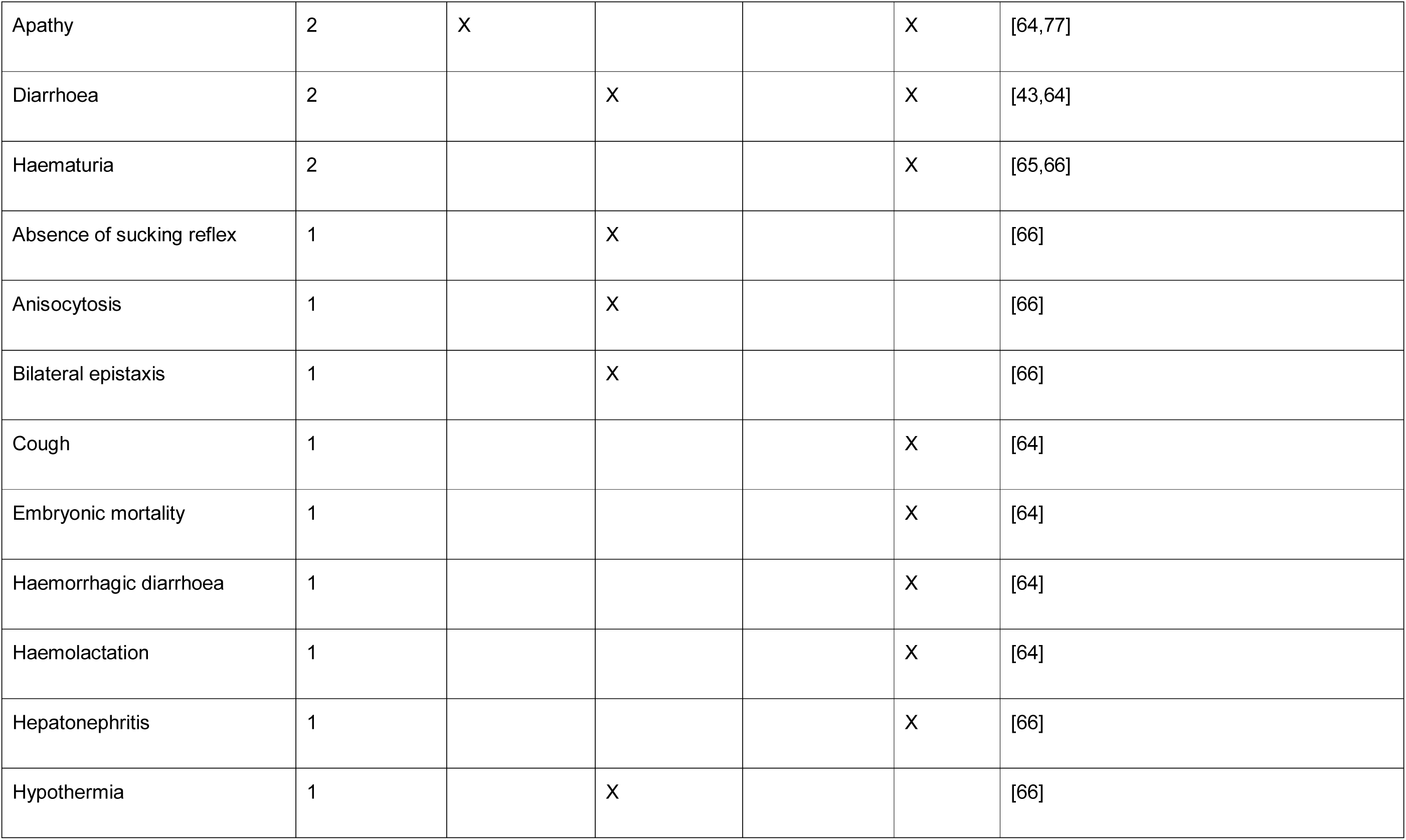
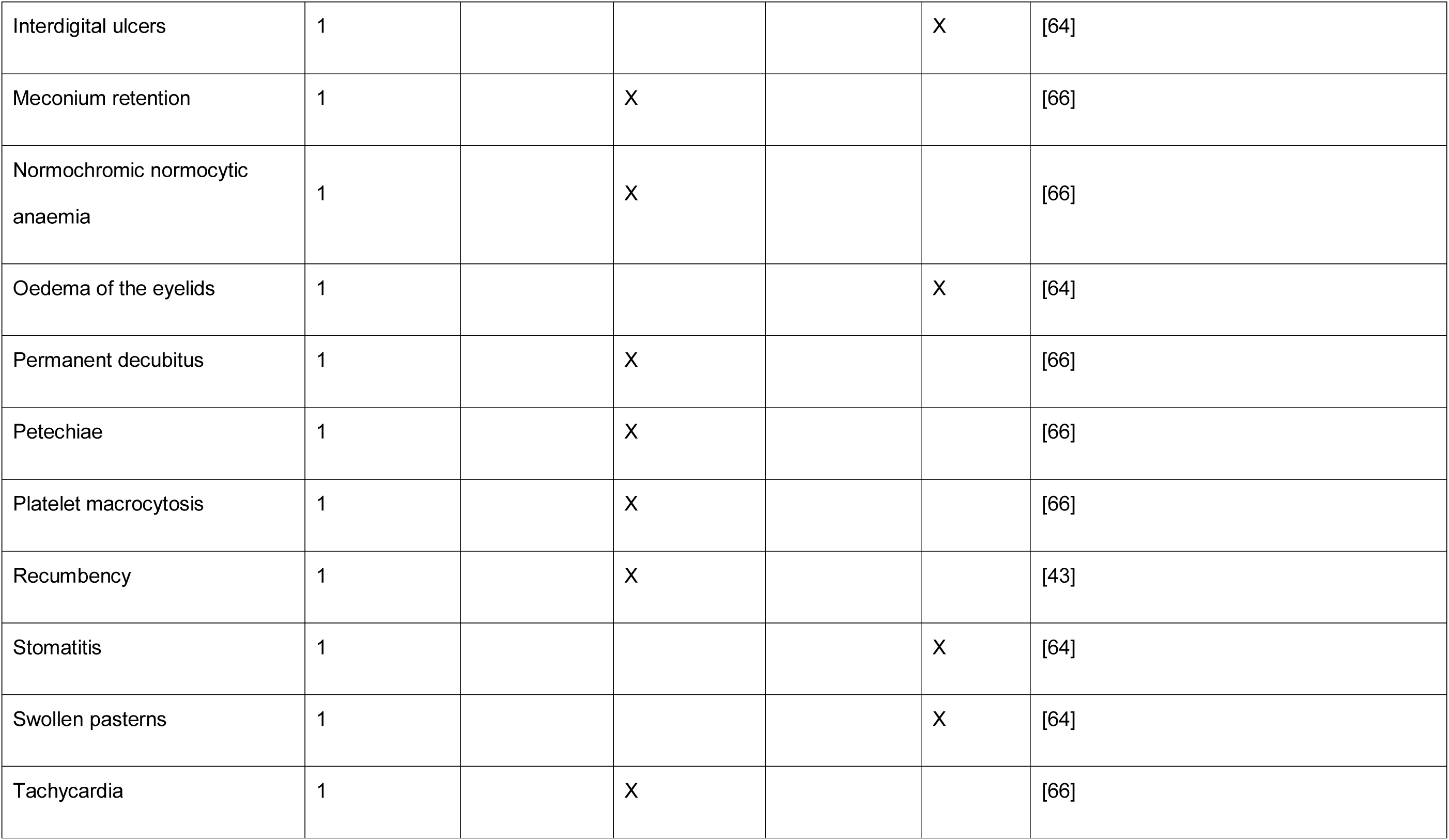
Clinical manifestations associated with bovine leptospirosis in Europe, 2001-2021. The table presents the number of studies reporting each sign age and the age category of the animals, as specified in the study (n = 16 studies).

| Clinical sign | Number of studies | Adult | Calf | Aborted fetus | Age not specified | References |
| --- | --- | --- | --- | --- | --- | --- |
| Abortion | 17 | X |  |  | X | [38–41,63,64,66,67,74,77,79,85,86,88,93,94,96] |
| Fertility disorders | 7 | X |  |  | X | [58,62,63,67,71,75,77] |
| Subclinical | 4 | X |  |  | X | [65,68–70] |
| Icterus | 4 |  | X | X |  | [43,66,85,86] |
| Perinatal mortality | 4 |  | X |  | X | [39,41,58,72] |
| Death | 3 |  | X |  | X | [42,64,66] |
| Decrease in milk production | 3 |  |  |  | X | [44,64,65] |
| Photosensitisation | 3 |  |  |  | X | [64–66] |
| Hyperthermia | 3 | X |  |  | X | [64,65,77] |
| Apathy | 2 | X |  |  | X | [64,77] |
| Diarrhoea | 2 |  | X |  | X | [43,64] |
| Haematuria | 2 |  |  |  | X | [65,66] |
| Absence of sucking reflex | 1 |  | X |  |  | [66] |
| Anisocytosis | 1 |  | X |  |  | [66] |
| Bilateral epistaxis | 1 |  | X |  |  | [66] |
| Cough | 1 |  |  |  | X | [64] |
| Embryonic mortality | 1 |  |  |  | X | [64] |
| Haemorrhagic diarrhoea | 1 |  |  |  | X | [64] |
| Haemolactation | 1 |  |  |  | X | [64] |
| Hepatonephritis | 1 |  |  |  | X | [66] |
| Hypothermia | 1 |  | X |  |  | [66] |
| Interdigital ulcers | 1 |  |  |  | X | [64] |
| Meconium retention | 1 |  | X |  |  | [66] |
| Normochromic normocytic anaemia | 1 |  | X |  |  | [66] |
| Oedema of the eyelids | 1 |  |  |  | X | [64] |
| Permanent decubitus | 1 |  | X |  |  | [66] |
| Petechiae | 1 |  | X |  |  | [66] |
| Platelet macrocytosis | 1 |  | X |  |  | [66] |
| Recumbency | 1 |  | X |  |  | [43] |
| Stomatitis | 1 |  |  |  | X | [64] |
| Swollen pasterns | 1 |  |  |  | X | [64] |
| Tachycardia | 1 |  | X |  |  | [66] |

Abortion was the most frequently reported sign of Leptospira infection in cattle (17/29, 58.6%) [38–41,63,64,66,67,74,77,79,85,86,88,93,94,96], followed by fertility disorders (7/29, 24.1%), including suboptimal reproductive performances such as prolonged calving intervals and poor conception rates [58,62,63,67,71,75,77]. These symptoms have been generally associated with chronic infections with Leptospira in European cattle [62,63,65,69,70,75], however, Grippi et al. mentioned infertility and abortion, associated with sudden death, in an acute episode of bovine leptospirosis [77]. A sudden drop in milk production was associated with an acute form of bovine leptospirosis in three studies [44,64,65]. Acute infections also manifested through hyperthermia [64,65,77], haemoglobinuria [65,66] and icterus [43,66,86]. In severe cases, death [42,64,66] and perinatal mortality [39,41,43,58,72] were described. Acute episodes in a cattle herd may be aggravated by concomitant infections, e.g. with Anaplasma phagocytophilum and bovine viral diarrhoea (BVD) virus [64].

Other clinical signs, e.g. photosensitisation [64–66], diarrhoea [43,64], haematuria [65,66], bilateral epistaxis [66], cough [66], swollen pasterns [64], interdigital ulcers [64] or oedema of the eyelids [64] were sporadically reported. Abnormalities in the blood count such as anisocytosis [66], platelet macrocytosis [66], or haemolytic anaemia [66] were also described.

Subclinical leptospirosis was reported in four studies [65,68–70]. Subclinical cases were more frequently found in dairy cattle than in beef herds, as well as abortion, fertility disorders, and hyperthermia. Death and apathy were described in both types of herds. Photosensitisation, perinatal mortality and drop in milk production were only reported in dairy herds. Systemic diseases, such as hypothermia, icterus, tachycardia, petechiae and changes in blood cell form (anisocytosis, platelet macrocytosis), were reported in beef production systems (Fig 6).

**Fig 6.**
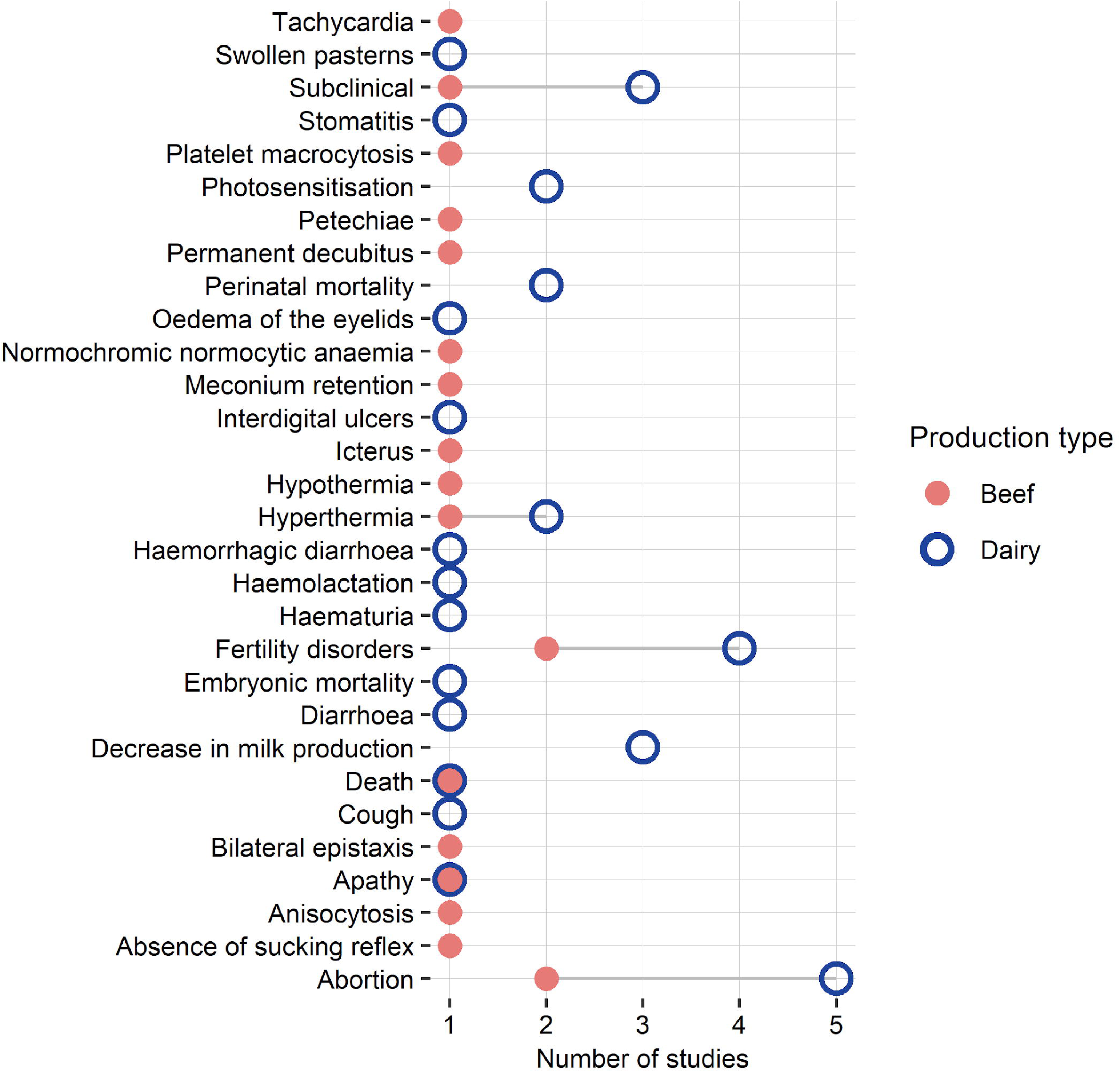
Cleveland dot plot displaying the number of studies reporting at least one clinical sign associated with bovine leptospirosis in Europe per type of production, 2001-2021.

Pathological findings were mostly described in Leptospira-positive calves and included pathological kidney alterations (4/5) [42,43,66,96], icterus [42,66,96], congested liver or hepatomegaly (3/5) [42,43,66], splenomegaly (3/5) [66,86,96] and petechiae on the lungs and heart (1/5) [66].

Histopathologically, changes in the liver, spleen, and kidney were most prominent [42,43,66,86,96]. Placentitis, necrosis, mononuclear or mixed inflammatory cell infiltrates, vasculitis and the presence of intracytoplasmatic and extracellular bacteria were observed on adult carcasses [88].

### Risk factors for bovine leptospirosis in Europe

Risk factors for bovine leptospirosis were addressed in 28/62 reviewed studies (45.2%) performed in 12 different countries [44,45,47,48,51–56,63–65,67,69–71,73,75,76,78,79,86,87,90,92,95,96]. Overall, a high diversity of risk factors and dependent variables were investigated (53 and 17, respectively, S1 Appendix, S2 Table). Environmental factors were the most commonly explored (n = 16 studies), followed by herd management practices (n = 10), biosecurity (n = 9), comorbidity with infectious diseases (n = 6), individual factors related to the animal (e.g. age, sex, breed) (n = 5), and clinical signs usually associated with leptospirosis (n = 4). Thirty-one risk factors were studied in dairy cattle while only 17 were investigated in beef herds, showing preferential investigations of dairy production systems (although production type was not mentioned in 10 studies). A statistical analysis, to quantify the relationship between the risk factor(s) and the dependent variable(s), was performed in 18/28 studies (64.3%); methods and models used were highly variable [45,47,48,51–56,67,70,71,73,76,78,86,87,96] (details in S1 Appendix). An overview of the statistically significant results is given in Fig 7, along with production type investigated and the number of studies investigating each risk factor (see also S1 Fig). Other studies evaluated qualitatively the risk through field observations [44,63–65,69,70,75,79,90,92,95].

**Fig 7.**
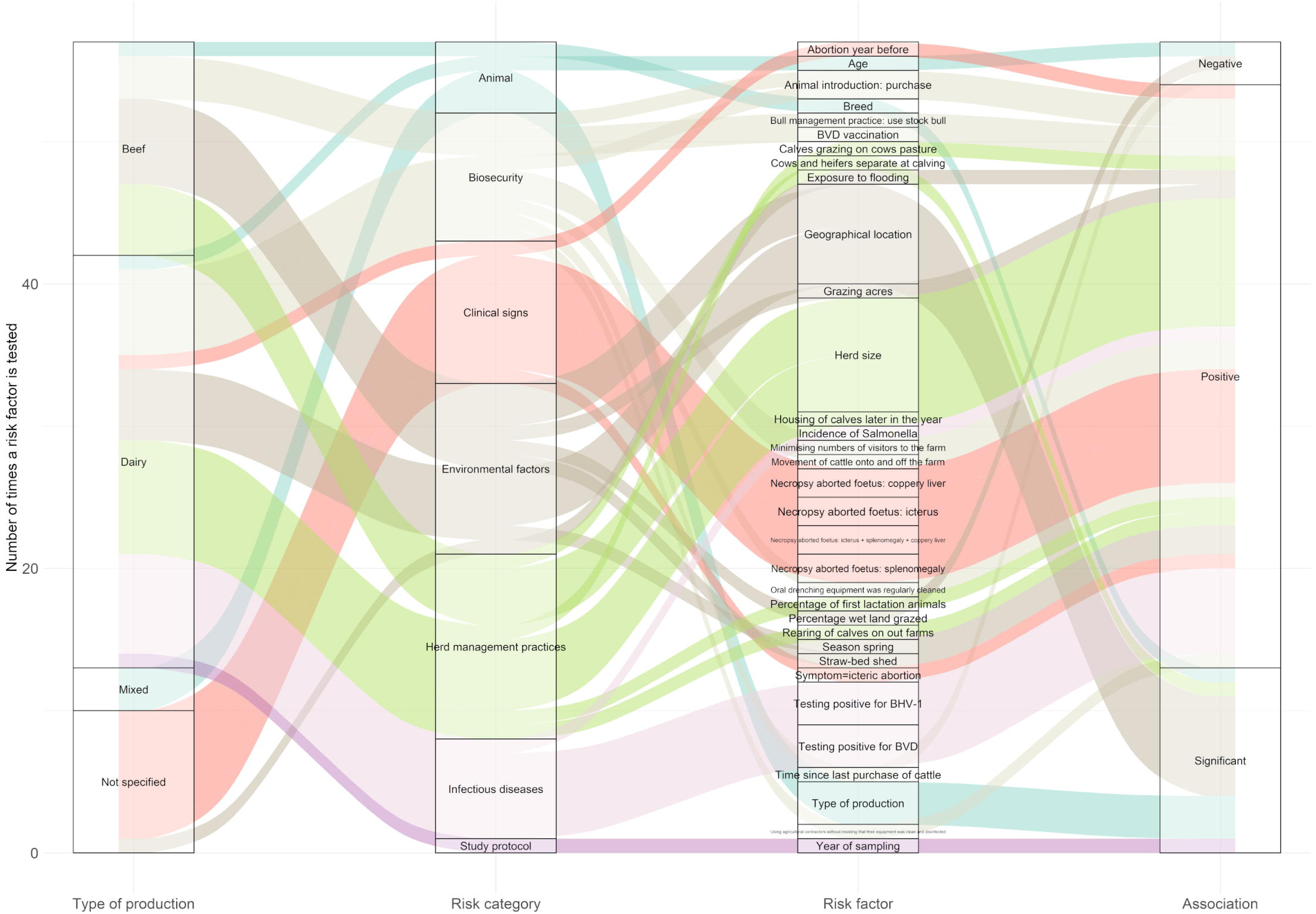
Sankey diagram showing the type of production, the category of the risk factor investigated, the risk factors investigated and the direction of the association with Leptospira-infection in cattle, Europe, 2001-2021. Association may be positive or negative; “significant” means that the association is statistically significant but no direction is provided. Only studies that have performed a statistical analysis (n = 18) are included and only significant risk factors are displayed. Colours represent the risk categories.

Five studies showed a significant statistical association between the geographic location of the animal (or herd) and Leptospira infection [48,52,55,56,76]. Flooding was significantly associated with higher antibody levels of Leptospira [87] while extreme weather conditions (i.e. warm/extremely warm or wet/extremely wet) were hypothetically (i.e. no statistical analysis was performed) associated with a high seroprevalence of leptospirosis [90]. In Spain, the season spring showed a significantly increased risk of infection with Leptospira serovar Grippotyphosa [71]. Access to pasture, grazing, and grazing surface area showed a significant positive association with Leptospira infection whereas increasing the percentage of wetland grazed significantly reduced the risk of infection [55]. The access to natural water sources might be a source of cattle infection (no statistical analysis performed) [44,75]. Moreover, the date of sampling exhibited a significant effect on Leptospira seroprevalence [76]. Certainly, date of sampling could be a confounding factor that may reflect the influence of e.g. the season, weather, or specific conditions at time of sampling.

The herd size was generally positively associated with an increased risk of Leptospira infection [45,47,51,52,54–56], although some studies found no statistical relationship [48,70,78]. Rearing calves off farm, co-grazing of calves and cows, housing the calves later in the year [54], or segregating heifers and cows at calving [55] significantly increased the risk of Leptospira infection. In addition, the percentage of primiparous cows in the herd was positively associated with the risk of herd infection [54]. The purchase of animals significantly increased the risk of Leptospira infection in animals and herds [47,51,69], although two studies did not confirm this result [45,55]. The movement of cattle onto and off the farm, e.g. for shows or temporary grazing [54], and the use of a stock bull [55] were significantly associated with the presence of Leptospira on farm. The employment of agricultural contractors untrained on biosecurity was evidenced as a risk factor of Leptospira infection in cattle whereas minimising the number of visitors was shown to decrease the probability of infection [54]. Surprisingly, one study reported that herds where cleaning of drenching equipment was performed were more likely to test positive for antibodies to Leptospira serovar Hardjo. The authors pointed out that one possible reason for the higher probability of a positive test for exposure to Leptospira is that cleaning of oral drenching equipment was carried out in response to the presence of the bacteria or other infectious diseases in the herd [54]. The presence of rodents (e.g. mice, nutria) [63,65] and the use of pig manure for the cow grazing pasture [75] were considered as risk factors of infection in cattle (although not tested statistically).

Past or co-occurring infections generally increased the risk of Leptospira infection while a significant positive association was reported between vaccination against the BVD virus and the seroprevalence of Leptospira [51]. Detection of circulating antibodies to BVD virus and bovine herpes virus 1 (BHV-1) increased the probability of detecting antibodies to Leptospira serovar Hardjo in the bulk milk [45,47,48]. Similarly, the incidence of Salmonella showed a significant statistical association with the incidence of Leptospira serovar Hardjo (such a link could not be proven for Neospora caninum) [53]. Finally, a previous Leptospira infection was not reported as a risk factor for subsequent infection [55] but recent history of abortion (i.e. the year before) was significantly associated with higher odds of testing seropositive for Leptospira serovar Bratislava [67]. Two studies demonstrated that icteric abortion was a strong predictor of Leptospira infection of the aborted dams [86] and foetuses [96]. Besides icterus, coppery liver and splenomegaly in the aborted foetuses were also significantly associated with infection by Leptospira [96].

Finally, animal characteristics such as age, sex, breed [56], or type of production [78] had an impact on Leptospira infection in cattle. Whereas age and sex of the cattle showed inconsistent effects on the risk of infection by Leptospira [56,73,75], dairy herds were significantly more at risk of infection by Leptospira serovar Copenhageni, Grippotyphosa, and Tarassovi than beef herds [78].

## Discussion

With 76 million head in 2021, Europe has a substantial cattle population [102]. To some extent, the geographic distribution of the publications included in this review reflects the distribution of the bovine population, with a large proportion of studies conducted in the United Kingdom, Ireland, and France. The annual number of published papers on cattle leptospirosis showed a positive trend over the study period which might indicate increasing level of interest in the disease in Europe. Despite the apparent growing risk, the impact on cattle health and welfare, and the potential severe economic losses related to bovine leptospirosis, there is a lack of comprehensive data on Leptospira infection in cattle in the European region, with studies published from 18 European countries only.

The occurrence of leptospirosis in European cattle varied greatly between countries and within countries, possibly depending on the geographical scale of the study (national, regional, or at farm level), specific location, the presence or absence of symptoms in the surveyed animals, the type of production system investigated (dairy versus beef), the study design (e.g. cross-sectional or longitudinal survey, case-control study), including the sample size which showed a large range, from one animal to more than 1 million cattle, the laboratory method used, and cut-off values considered. Knowledge of the (sero)prevalence of bovine leptospirosis in a geographic region is essential for veterinary practitioners to include the disease in their differential diagnoses and reduce underdiagnoses and misdiagnoses. Many studies reported seroprevalences based on a sample size < 50 cattle and/or were conducted in a limited geographic area (sometimes at farm level), probably because clinical case reports represented one quarter of the included studies. Consequently, the true prevalence of the bacteria in Europe remains elusive.

In the examined studies, all attempts to culture Leptospira from field samples failed. Europe, and especially bovine species, are underrepresented in the Leptospira isolates and genomes database of the Institut Pasteur, which includes only nine isolates from cattle/bovine in Europe (out of 153 European isolates from animals, humans, and the environment, i.e. 5.9%) [103], while the database includes a total of 83 Leptospira strains isolated from cattle/bovines worldwide (out of 745 deposited isolates, 11.1%). Isolation of local strains from infected animals, humans, or the environment is essential to improve the sensitivity of the MAT for diagnosis in humans and animals, as well as to further evaluate the genetic diversity of the genus at local, regional, national, and international scale [104,105]. The recovery of Leptospira from field samples is extremely challenging, however isolated strains are indispensable to phylogenetic analyses.

This review exposed the heterogeneity of the methods used to confirm leptospirosis in cattle in Europe; these methods certainly exhibit variable inter-laboratory performances. Likewise, the cut-off values proposed for the MAT displayed significant variations, with a lack of consensus over which titre should be used for a positive result. Interpretation of the MAT titre should consider the vaccination history of the animals tested, since the MAT and ELISA do not differentiate between natural and vaccine-induced antibodies [104], as well as the epidemiological context, since limitations in the diagnosis are reported in case of chronic infection and in herds (or regions) where the disease is endemic [104]. In addition, cut-off values to be considered during a clinical case investigation versus an epidemiological survey should be different. Also, the panel of serovars used for the MAT was found to be extremely variable between studies, which prevents accurately mapping serovar/serogroup distribution in Europe. Lastly, intrinsic limitations of the method, i.e. cross-reactivity between serovars in the MAT, does not allow the identification of the infecting serovar with absolute certitude [104]. Moreover, very often, the agglutinating serovars with the highest MAT titres is the only one reported, narrowing our perspective on the epidemiology of the circulating serovars/serogroups.

The World Organisation for Animal Health (WOAH) recommends that the MAT panel used for serodiagnosis of leptospirosis in animals includes representative strains of the serogroups known to circulate locally as well as those known to circulate in the animal species elsewhere [104]. This review demonstrated that the serogroups Sejroe (for which cattle are recognised as maintenance host), Australis, Grippotyphosa, Icterohaemorrhagiae, and Pomona are the most reported ones in cattle in Europe and should therefore be included in the basic MAT panel for the serodiagnosis of leptospirosis in cattle in this region. This finding supports previous observations, e.g. in South America [34,106,107] Africa [108,109], Malaysia [110], New Zealand [111] and Australia [112], where serogroup Sejroe is predominantly reported in bovines. Serogroups Australis, Grippotyphosa, Icterohaemorrhagiae and Pomona have also been described in cattle worldwide [34,106–109,111,113,114]. Nevertheless, those serogroups are the most used in the MAT panels, which very likely biases the overall picture.

Differences in the number and type (e.g. clinical case report and case-control studies versus transversal survey) of studies in the different countries may induce a bias in the reported clinical signs. For example, abortion was mainly reported in studies from the United Kingdom, Belgium and France, which counted a high number of published articles but also a greater proportion of case reports or case-control studies on abortion problems. It is therefore not possible to decipher from the data if there is a geographic difference in the clinical expression of the disease (which might be caused, for example, by differences in the infecting serovars [115]) or if there is, perhaps, an inherent researcher bias on reporting or investigating certain clinical signs in some countries. Overall, this work highlights that the clinical features of bovine leptospirosis in Europe are diverse, covering a wide range of symptoms, making a field diagnosis challenging. The diagnosis should therefore rely on laboratory tests. As described in tropical and sub-tropical areas [116,117], New Zealand [118], and Australia [119], the most recognised and reported clinical signs associated with bovine leptospirosis in European cattle are related to suboptimal reproductive performance, such as abortions, fertility disorders, stillbirths, birth of weak offspring, poor conception rate and prolonged calving intervals. Therefore, in case of abortion events, leptospirosis should be considered as a differential diagnosis among other infectious causes of abortion, i.e. Brucella spp., Neospora caninum, Coxiella burnetii, BVD virus, BHV-1, or Salmonella enterica serotype Dublin [120,121].

This review provides a comprehensive overview of the risk factors of Leptospira infection in cattle in Europe. However, the heterogeneity of approaches, i.e. regarding study design, diagnostic method, statistical model, dependent and explanatory variables, sample size, and epidemiological unit, challenges the interpretation and comparison of the results. In general, the most reported significant risk factors of leptospirosis were related to biosecurity measures, e.g. livestock purchase and quarantine policy, rodent control, separation of age groups, mixing with other species, or use of a stock bull. Environmental context plays also a major role in the risk of infection, e.g. geographic location. Local or even regional environmental conditions influence the presence of Leptospira in the environment as well as host-bacteria interactions. Furthermore, several factors related to the soil and water pH, temperature, or composition of the environmental microbiome may determine the possibility of persistence of Leptospira outside its animal host [122–124]. Climatic conditions in Europe are becoming increasingly suitable for the survival and transmission of water-and rodent-borne diseases, including leptospirosis [125]. Extreme weather events compounding the impact of changes in land use (especially urbanisation) intensify the direct and indirect (i.e. via the environment) contacts between leptospires, humans, maintenance and susceptible animals [4,126–128], therefore increasing the risk for public and animal health. Finally, this research shows that concomitant infections (e.g. BVD, BHV-1, Neospora caninum, Salmonella, etc.) increase the risk for Leptospira infection. This hypothesis is supported by studies from Laos, where seropositivity to Leptospira was associated with higher titres of N. caninum and BVD virus [129]. We can hypothesise that the lack of biosecurity measures and hygiene on infected farms favours not only infection by Leptospira, but also by other pathogens. Furthermore, animals infected with other diseases are likely to experience a certain level of immunosuppression and be more susceptible to infection with Leptospira [130–132]. Another hypothesis is that farms infected with multiple pathogens have less frequent veterinary visits; for example, in Ecuador and Brazil, farms with no or limited veterinary services have a higher risk of infection with Leptospira [133,134]. Overall, this review demonstrated the multiplicity and complexity of a variety of entangled risk factors associated with Leptospira exposure in cattle. These findings should motivate local and national health authorities, veterinarians, and farmers to implement integrated disease prevention and control measures at farm or regional level.

Research on cattle leptospirosis in Europe suffers several data and research gaps. Notably, this review highlights shortcomings in the diagnostic methods. Harmonised protocols to investigate, diagnose, and report cases of leptospirosis in cattle are necessary. Cattle-specific cut-off values for the MAT should be defined and based on the epidemiological context, e.g. endemic versus non-endemic region, vaccinated versus non-vaccinated herd. To provide comparable data at European level, a pan-European consensus is needed regarding the minimum panel of strains to be used for the MAT, which could further be regionally optimised with locally-isolated strains [135]. Moreover, the isolation and molecular characterisation of Leptospira from cattle will advance knowledge on the epidemiology, ecology, and pathogenesis of the bacteria, and will have practical applications in the prevention, surveillance, and control of the disease in both animals and humans. More efforts are needed in this direction. This review also stresses a lack of large-scale studies, necessary for drawing representative conclusions and achieving sufficient statistical power and points out that dairy herds are disproportionally more frequently investigated compared to beef herds, leading to a data gap regarding the clinical signs, prevalence, and impact of the disease in beef cattle. Additionally, we noted a lack of data on the risk related to artificial insemination and a limited investigation of rodents as a source of Leptospira infection in cattle herds. Finally, in the last 20 years, no study investigated bovine genital leptospirosis (BGL) in Europe, although studies from Brazil have evidenced a relatively high prevalence of the disease [13,136,137] and point toward the recognition of BGL as a distinct syndrome [138].

## Conclusions

Research on bovine leptospirosis is generally under-resourced while the disease is globally neglected [139], including in Europe, where a limited number of countries have investigated and reported the disease. Considering the veterinary and public health importance of leptospirosis as well as its economic impact, it is crucial to raise awareness among stakeholders, including farmers, veterinarians, and other health professionals, in areas where the disease is not (yet) endemic. This is especially important in Europe where this zoonotic disease is (re-)emerging in humans, but also in animals. Moreover, intensification of livestock farming in certain regions of Europe, concomitant with an increasing trend toward herd grazing outdoors in other areas will also certainly play a major role in the future regional incidences of leptospirosis in cattle. Local studies are essential to advance our understanding of the epidemiology of bovine leptospirosis and therefore develop and implement relevant, locally-adapted prevention and control strategies. Nevertheless, an overview of the epidemiology of the disease at continental scale, as presented here, can yield novel insights into its epidemiological features, especially, by unfolding common determinants of disease events.

## Supporting information

S1 PRISMA Checklist

## Supporting information

**S1 Appendix. Extracted data on cattle leptospirosis retrieved from the 62 reviewed papers, Europe, 2001-2021.** Sheet 1: Extracted data (except risk factors); Sheet 2: Extracted data on risk factors of cattle leptospirosis in Europe; Sheet 3: Risk categories attributed to the different risk factors; Sheet 4: Metadata.

**S1 Table. Search queries used in the different databases for this systematic literature review.**

**S2 Table. Dependent variables pertaining to infection with Leptospira spp. in cattle and number of studies that used them to decipher risk factors of infection, Europe, 2001-2021.**

**S1 Fig. Forest plot of the risk factors of bovine leptospirosis, Europe, 2001-2021.** The figure presents the odds ratios (OR) and 95% confidence interval (95% CI) for nine European studies that used this metrics to evaluate risk factors of leptospirosis in cattle. Note that if 95% CI contains the value 1, the OR is not significant.

**S1 PRISMA Checklist. Checklist for reporting systematic reviews.**

## Notes

### Competing Interest Statement

The authors have declared no competing interest.

