## Supplementary material for "A systematic review on leptospirosis in cattle: a European perspective": S1 PRISMA Checklist

### Reporting checklist for systematic reviews.

Based on the PRISMA guidelines.

| | | Reporting Item | Page Number |
|---|---|---|---|
| **Title** | | | |
| Title | #1 | Identify the report as a systematic review | 1 |
| **Abstract** | | | |
| Abstract | #2 | Report an abstract addressing each item in the PRISMA 2020 for Abstracts checklist | 2 |
| **Introduction** | | | |
| Background/rationale | #3 | Describe the rationale for the review in the context of existing knowledge | 4-5 |
| Objectives | #4 | Provide an explicit statement of the objective(s) or question(s) the review addresses | 5-6 |
| **Methods** | | | |
| Eligibility criteria | #5 | Specify the inclusion and exclusion criteria for the review and how studies were grouped for the syntheses | 6-7 |
| Information sources | #6 | Specify all databases, registers, websites, organisations, reference lists, and other sources searched or consulted to identify studies. Specify the date when each source was last searched or consulted | 6 |
| Search strategy | #7 | Present the full search strategies for all databases, registers, and websites, including any filters and limits used | 6, S1 Table |
| Selection process | #8 | Specify the methods used to decide whether a study met the inclusion criteria of the review, including how many reviewers screened each record and each report retrieved, whether they worked independently, and, if applicable, details of automation tools used in the process | 6-7 |
| Data collection process | #9 | Specify the methods used to collect data from reports, including how many reviewers collected data from each report, whether they worked independently, any processes for obtaining or confirming data from study investigators, and, if applicable, details of automation tools used in the process | 7 |
| Data items | #10a | List and define all outcomes for which data were sought. Specify whether all results that were compatible with each outcome domain in each study were sought (for example, for all measures, time points, analyses), and, if not, the methods used to decide which results to collect | 7 |
| Data items | #10b | List and define all other variables for which data were sought (such as participant and intervention characteristics, funding sources). Describe any assumptions made about any missing or unclear information | 7 |
| Study risk of bias assessment | #11 | Specify the methods used to assess risk of bias in the included studies, including details of the tool(s) used, how many reviewers assessed each study and whether they worked independently, and, if applicable, details of automation tools used in the process | n/a |
| Effect measures | #12 | Specify for each outcome the effect measure(s) (such as risk ratio, mean difference) used in the synthesis or presentation of results | 7 |
| Synthesis methods | #13a | Describe the processes used to decide which studies were eligible for each synthesis (such as tabulating the study intervention characteristics and comparing against the planned groups for each synthesis (item #5)) | 7 |
| Synthesis methods | #13b | Describe any methods required to prepare the data for presentation or synthesis, such as handling of missing summary statistics or data conversions | 7 |
| Synthesis methods | #13c | Describe any methods used to tabulate or visually display results of individual studies and syntheses | 7 |

| | | | |
|---|---|---|---|
| Synthesis methods | #13d | Describe any methods used to synthesise results and provide a rationale for the choice(s). If meta-analysis was performed, describe the model(s), method(s) to identify the presence and extent of statistical heterogeneity, and software package(s) used | S1 Appendix |
| Synthesis methods | #13e | Describe any methods used to explore possible causes of heterogeneity among study results (such as subgroup analysis, meta-regression) | n/a |
| Synthesis methods | #13f | Describe any sensitivity analyses conducted to assess robustness of the synthesised results | n/a |
| Reporting bias assessment | #14 | Describe any methods used to assess risk of bias due to missing results in a synthesis (arising from reporting biases) | n/a |
| Certainty assessment | #15 | Describe any methods used to assess certainty (or confidence) in the body of evidence for an outcome | n/a |
| **Results** | | | |
| Study selection | #16a | Describe the results of the search and selection process, from the number of records identified in the search to the number of studies included in the review, ideally using a flow diagram (http://www.prisma-statement.org/PRISMAStatement/FlowDiagram) | 8, Fig1 |
| Study selection | #16b | Cite studies that might appear to meet the inclusion criteria, but which were excluded, and explain why they were excluded | n/a |
| Study characteristics | #17 | Cite each included study and present its characteristics | S1 Appendix, Fig1, Fig2 |
| Risk of bias in studies | #18 | Present assessments of risk of bias for each included study | n/a |
| Results of individual studies | #19 | For all outcomes, present for each study (a) summary statistics for each group (where appropriate) and (b) an effect estimate and its precision (such as confidence/credible interval), ideally using structured tables or plots | S1 Appendix, S1Fig |
| Results of syntheses | #20a | For each synthesis, briefly summarise the characteristics and risk of bias among contributing studies | 27-28 |
| Results of syntheses | #20b | Present results of all statistical syntheses conducted. If meta-analysis was done, present for each the summary estimate and its precision (such as confidence/credible interval) and measures of statistical heterogeneity. If comparing groups, describe the direction of the effect | Table 1, Table 2, Table 3 Fig5, Fig6, Fig7 |
| Results of syntheses | #20c | Present results of all investigations of possible causes of heterogeneity among study results | 27-28 |
| Results of syntheses | #20d | Present results of all sensitivity analyses conducted to assess the robustness of the synthesised results | n/a |
| Risk of reporting biases in syntheses | #21 | Present assessments of risk of bias due to missing results (arising from reporting biases) for each synthesis assessed | n/a |
| Certainty of evidence | #22 | Present assessments of certainty (or confidence) in the body of evidence for each outcome assessed | n/a |
| **Discussion** | | | |
| Results in context | #23a | Provide a general interpretation of the results in the context of other evidence | 27 |
| Limitations of included studies | #23b | Discuss any limitations of the evidence included in the review | 30 |
| Limitations of the review methods | #23c | Discuss any limitations of the review processes used | 29 |
| Implications | #23d | Discuss implications of the results for practice, policy, and future research | 31 |
| **Other information** | | | |
| Registration and protocol | #24a | Provide registration information for the review, including register name and registration number, or state that the review was not registered | n/a |
| Registration and protocol | #24b | Indicate where the review protocol can be accessed, or state that a protocol was not prepared | n/a |
| Registration and protocol | #24c | Describe and explain any amendments to information provided at registration or in the protocol | n/a |

| | | | |
|---|---|---|---|
| Support | #25 | Describe sources of financial or non-financial support for the review, and the role of the funders or sponsors in the review | Financial disclosure (online submission) |
| Competing interests | #26 | Declare any competing interests of review authors | Competing interests (online submission) |
| Availability of data, code, and other materials | #27 | Report which of the following are publicly available and where they can be found: template data collection forms; data extracted from included studies; data used for all analyses; analytic code; any other materials used in the review | S1 Appendix |

The PRISMA checklist is distributed under the terms of the Creative Commons Attribution License CC-BY. This checklist can be completed online using https://www.goodreports.org/, a tool made by the EQUATOR Network in collaboration with Penelope.ai
