## Supplementary material for "A systematic review on leptospirosis in cattle: a European perspective": S1 Table

**S1 Table. Search queries used in the different databases for this systematic literature review.**

| Database | Search query |
|---|---|
| **Pubmed** | (((lepto*) AND ((cattle) OR (cow) OR (cows)) |
| **Scopus** | TITLE-ABS-KEY ( lepto* ) AND TITLE-ABS-KEY ( cattle OR cow OR cows ) |
| **Web of Science** | (((AB=(lepto*)) AND (AB=(cattle) OR AB=(cow) OR AB=(cows)))) |
| **CABI** | ((lepto*) AND ((cattle) OR (cow) OR (cows)) AND yr:[2001 TO 2021]) AND (((geographic-location:(( "Italy" OR "Poland" OR "Irish Republic" OR "Germany" OR "Spain" OR "Russia" OR "Wales" OR "Netherlands" OR "Lombardy" OR "Austria" OR "Bulgaria" OR "UK" OR "England" OR "Croatia" OR "Nordic Countries" OR "Czech Republic" OR "Bosnia-Hercegovina" OR "Belgium" OR "Switzerland" OR "Northern Ireland" OR "Slovakia" OR "Europe" OR "France" ) )) (language:(( "English" OR "German" OR "French" ) )) )) |
