## Supplementary material for "A systematic review on leptospirosis in cattle: a European perspective": S2 Table

**S2 Table. Dependent variables used in the studies investigating risk factors of cattle leptospirosis in Europe, 2001-2021 (n = 28 studies).** sv.: serovar; sg.: serogroup.

| **Dependent variable** | **Number of studies** | **References** |
|---|---|---|
| Herd (sero)positivity to *Leptospira* | 9 | [45,47,48,52-55,79,87] |
| Within-herd seroprevalence | 7 | [45,47,48,51,56,70,78] |
| Seroprevalence | 6 | [56,73,76,90,92,95] |
| Animal (sero)positivity to *Leptospira* | 5 | [44,63,65,75,86] |
| Incidence of *Leptospira* | 2 | [53,69] |
| Animal seropositivity to *Leptospira* (sv. Bratislava) | 1 | [67] |
| Herd antibody titre level | 1 | [87] |
| Herd seroprevalence | 1 | [56] |
| Incidence of *Leptospira* sv. Grippotyphosa | 1 | [71] |
| Incidence of *Leptospira* sg. Australis (sv. Bratislava) | 1 | [71] |
| PCR positive in aborted foetus | 1 | [96] |
| Seropositivity to *Leptospira* in aborted dam (titre 1:100) | 1 | [96] |
| Severity of clinical symptoms | 1 | [64] |
| Within-herd antibody titre level | 1 | [52] |
| Within-herd seroprevalence (sv. Copenhageni) | 1 | [78] |
| Within-herd seroprevalence (sv. Grippotyphosa) | 1 | [78] |
| Within-herd seroprevalence (sv Tarassovi) | 1 | [78] |
