## Supplementary figures and images for "A systematic review on leptospirosis in cattle: a European perspective"

### S1 Figure

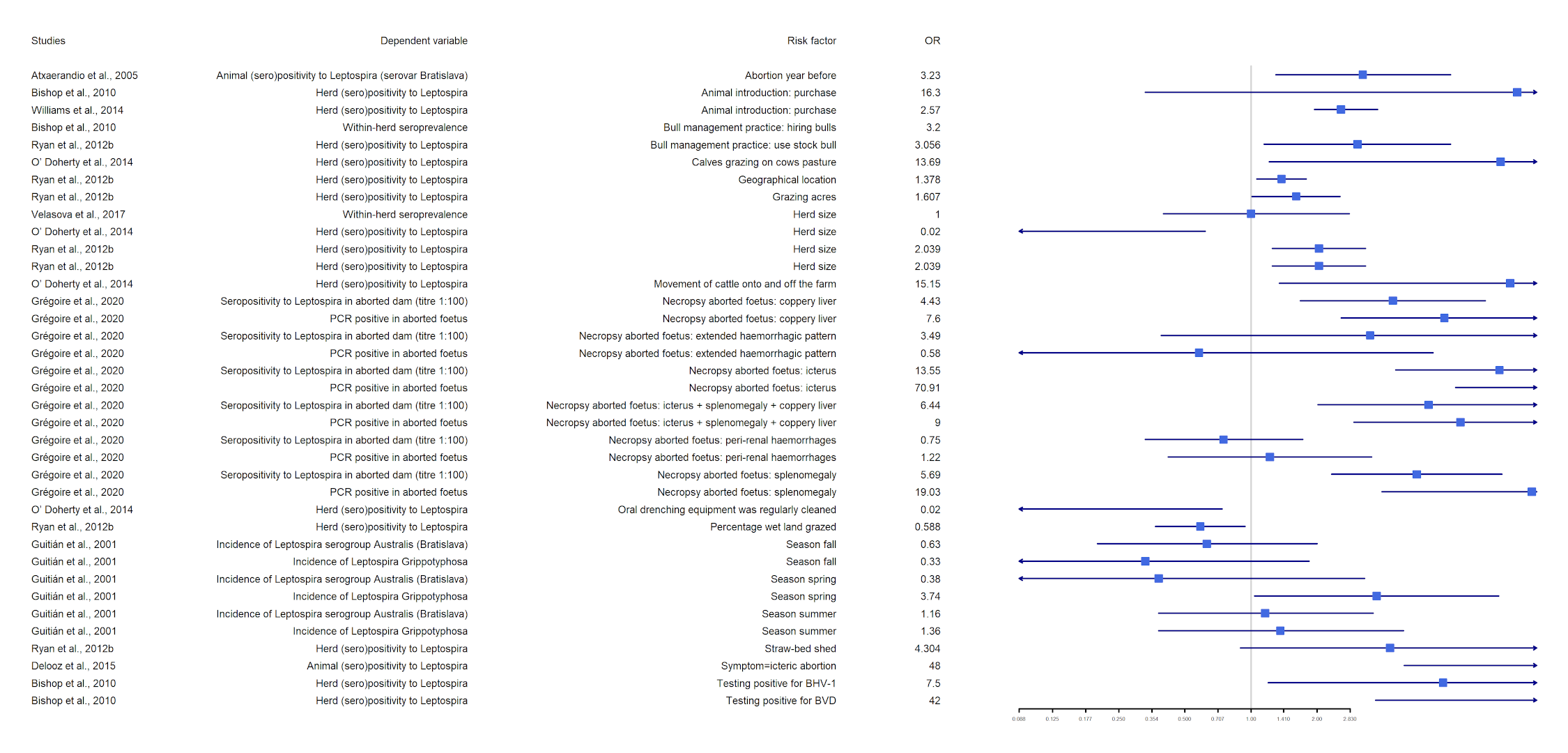
